# Quantitative microbiology with widefield microscopy: navigating optical artefacts for accurate interpretations

**DOI:** 10.1101/2023.05.15.540883

**Authors:** Georgeos Hardo, Ruizhe Li, Somenath Bakshi

## Abstract

Time-resolved live-cell imaging using widefield microscopy is instrumental in quantitative microbiology research. It allows us to track and measure the size, shape, and content of individual microbial cells over time. However, the small size of microbial cells poses a significant challenge in interpreting image data, as it approaches the dimensions of the microscope’s depth of field and experiences diffraction effects. As a result, 2D widefield images of microbial cells contain projected 3D information, blurred by the 3D point spread function. In this study, we employ computer simulations and targeted experiments to investigate the impact of diffraction and projection on our ability to quantify the size and content of microbial cells from 2D microscopic images. This study points to some new and often unconsidered artefacts resulting from the interplay of projection and diffraction effects, within the context of quantitative microbiology. These artefacts introduce substantial errors and biases in size and fluorescence quantification, and even single-molecule counting, making the elimination of these errors a complex task. Awareness of these artefacts is crucial for designing strategies to accurately interpret micrographs of microbes. To address this, we present new experimental designs and machine learning based analysis methods that account for these effects, resulting in accurate quantification of microbiological processes.

---

Widefield fluorescence microscopy is a cornerstone of quantitative microbiology, allowing for noninvasive, real-time imaging of individual cells. This technique’s capacity to measure the size, shape, and content of individual microbial cells has advanced several areas of quantitative microbiology research, including studies on size regulation and division control in bacteria^1–3^, regulation and noise in gene expression^4–6^, and analysis of interactions between cells and their environments^7,8^. Additionally, imaging individual molecules within cells using this technique has enabled the study of the dynamics and organisation of individual genes, mRNAs, and proteins, and has facilitated the construction of accurate distributions of their abundance^9,10^. In essence, live-cell widefield microscopy plays a pivotal role in developing a comprehensive understanding of biological processes within and between microbial cells, offering insights into their organisation, dynamics, and regulation across different scales.

However, careful scrutiny is required for the extraction of quantitative microbiological information from microscopy data. The size of most microbes (particularly bacterial cells) is comparable to the dimensions of the microscope’s point-spread function (PSF)^11^, resulting in significant diffraction effects on bacterial cell images (blur). Moreover, the thickness of bacterial cells is roughly equivalent to the depth of field (DoF) of a microscope objective. Consequently, 2D widefield images of microbes contain projected 3D information containing diffraction from the 3D PSF. The interplay of projection and diffraction effects can bias the estimation of cell size, shape, and intensity. Additionally, these factors hinder the quantification of low copy-number molecules like mRNAs and transcription factors from single-molecule counting experiments^10^ as molecules at varying depths exhibit varying degrees of defocus, overlap in the 2D projection, and coalesce into a single blurred spot due to diffraction. Systematically analysing these effects to understand their impact on accurate interpretation of microscopy data has proven challenging due to the lack of accurate ground-truth information.

To tackle this challenge, we utilised SyMBac (Synthetic Micrographs of Bacteria), a virtual microscopy platform (which we introduced in a previous work^12^) capable of generating synthetic bacterial images under various conditions. This tool allows us to assess diffraction and projection effects through forward simulation. The SyMBac-generated images come with precise ground-truth data, enabling us to accurately quantify errors and biases in different measurements and offer control over a wide range of parameters, encompassing optics, physics, and cell characteristics (size, shape, intensity distribution, and fluorescent label type). Consequently, we can analyse how these factors affect image formation and feature extraction. Moreover, the virtual microscopy setup allows us to explore imaging parameters that may be difficult or impractical to realise in actual experiments but are crucial for identifying important variables by amplifying their effects.

In this paper, we use SyMBac to systematically investigate the impact of projection and diffraction on the accurate quantification of three key aspects: 1. Cell dimensions, 2. Fluorescence intensity of individual cells, and 3. Counts of individually labelled entities per cell. To validate the findings from our virtual microscopy experiments, we conducted targeted real experiments with variable optical settings. Our analysis revealed previously unrecognised artefacts arising from the interplay of projection and diffraction effects. These artefacts introduce significant errors and biases in the estimation of cell size, intensity, and molecule counts, proving challenging to rectify. Recognizing these effects and devising appropriate mitigation strategies is crucial for accurate quantification of microbiological processes from microscopy data. To this end, we have demonstrated that understanding these effects enables designing ‘smart-microscopy’ experiments, along with analytical protocols that minimise their impact while facilitating accurate data interpretation for estimating cell size and content measurements.

## Results

### Digital widefield fluorescence microscopy experiments

We employed the SyMBac virtual microscopy platform to conduct digital experiments mimicking widefield epifluorescence microscopy, the technique typically used for time-resolved live-cell microbial imaging^13,14^. In this configuration, microbial cells are sandwiched between a glass coverslip and a biocompatible material like agarose or PDMS, as they are imaged through the cover glass via the microscope’s objective lens, which can be either upright or inverted (Fig. 1a).

**Figure 1:**
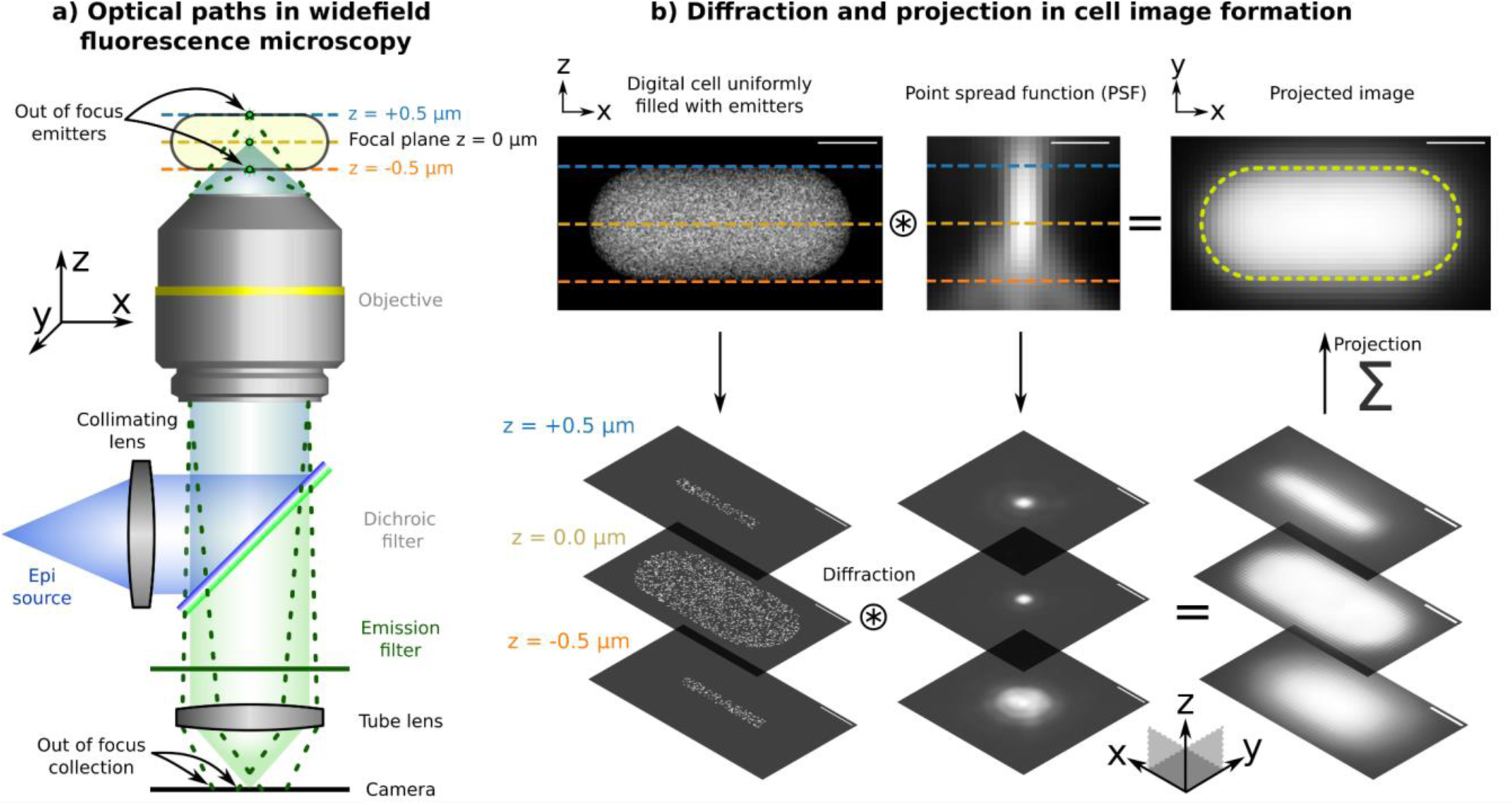
Projection and diffraction effects in widefield fluorescence image formation: **a)** Schematic optical paths for sample illumination (blue) and emission (green) collection on the camera for image formation in an epi-fluorescence setup. Emitters in the midplane of the cell are in focus. **b)** A stepwise illustration of image formation of a cell uniformly filled with fluorescent emitters. Light from emitters at various planes of the cell is diffracted by the corresponding plane of the 3D point spread function (PSF). Images from multiple planes at various sample depths are projected on top of each other to form the final 2D image of the cell. Further details on the image formation process are given in Supplementary Information 1. All scale bars are 0.5 μm.

In these settings, the microscope objective has a dual role: it focuses the excitation beam, illuminating the sample through the cover glass and collects emitted photons from the entire sample, focusing them on the camera to form the image (Fig. 1a). The excitation light illuminates the entire sample, inducing fluorescence throughout. The objective collects emitted light from various planes along the sample’s Z-axis within its depth of field. Each of these planes in the Z-axis introduces blurring due to the 3D PSF, resulting in the projected 2D image comprising contributions from each Z-plane. Each contribution is differentially blurred according to its relative distance from the focal plane and the corresponding slice of the 3D PSF (Fig. 1b). The interplay of projection and diffraction effects in image formation presents significant challenges in accurately extracting the ground-truth distribution of the emitters, as elaborated below.

### Effects of projection and diffraction on cell size estimation

We start by examining how projection and diffraction impact the quantification of cell size and shape from 2D fluorescence images. The extent of blurring due to “diffraction effects” is linked to the 3D PSF’s size, which depends on imaging wavelength, the numerical aperture of the objective lens, and any aberrations within the optical system. Consequently, diffraction effects exhibit wavelength-dependent characteristics for a fixed objective lens. In our digital simulations and real experiments, we employed PSFs of different wavelengths to investigate how diffraction impacts error and bias in measurements. Conversely, the manipulation of the depth of field in the imaging setup reveals the influence of “projection effects.” Using SyMBac, we can selectively toggle projection or diffraction effects, thus allowing us to model each effect in isolation by either capturing light from an infinitesimally thin plane or omitting convolution with the PSF, as shown in Fig. 2a and g. These “nonphysical” experiments are instrumental in identifying and understanding each underlying effect and its contribution.

**Figure 2-.**
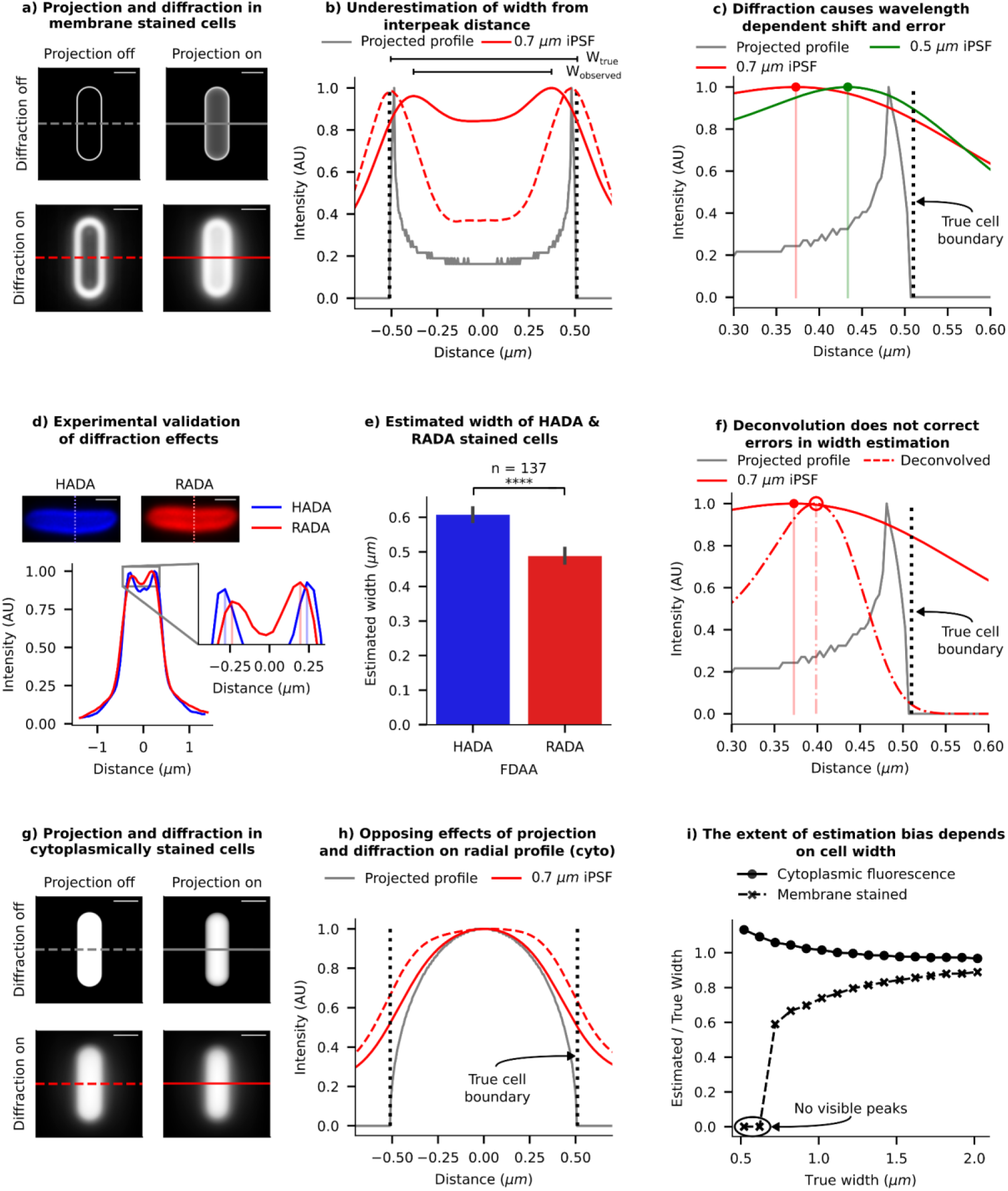
Effects of projection and diffraction on cell size estimation: **a)** Synthetic images of a stained cell membrane demonstrate the independent and combined effects of diffraction and projection on 2D image formation (scale bar = 1 µm). Diffraction effects were simulated using the experimentally measured instrumental PSF (iPSF) of our imaging system. **b)** Radial profiles of intensity (across the cell width) from each panel from a) are compared to show the relative shifts caused by projection and diffraction effects. The black dotted line indicates the true position of the cell boundary. **c)** Radial profiles of synthetic-cell images with two wavelengths of iPSFs are shown. Lower wavelengths (green) cause smaller shifts, as expected from a smaller size of the PSF. **d)** Example images of a cell stained with fluorescent D-amino acids HADA and RADA are shown. Intensity traces across the cell’s width show RADA (red) emission is more diffracted than HADA (blue) emission, and the diffraction is biassed towards the centre of the cell. **e)** A plot showing the average measured width of a population of cells stained with HADA and RADA (error bar = 99% CI). Inter-peak distances from radial profiles of RADA images consistently underestimate the width more than HADA images. **f)** Comparison of radial profiles before and after deconvolution shows that deconvolution does not shift and correct the peak position; it only makes the profile sharper. **g)** Synthetic images of a digital cell, uniformly filled with fluorescence emitters, show the effects of diffraction and projection on 2D image formation (scale bar = 1 µm). **h)** We compare the radial profile of intensity (across the cell width) with and without projection and diffraction effects corresponding to the panels in **g)**. The black dotted line indicates the true position of the cell boundary. **i)** Trendlines from synthetic data show that the observed/true width ratio is dependent on the cell width, with the error growing rapidly for narrow cells. The trend, however, occurs in opposite directions for membrane stained cells and cytoplasm-labelled cells. Estimated widths are calculated from interpeak distance in membrane-stained cells and full width at half maximum (FWHM) of the radial profile of cytoplasm-labelled cells.

### Errors and biases in size estimation from membrane-stained images

To define cell boundaries, quantitative microbiologists often use a membrane stain, a fluorescent dye that highlights the cell membrane (or the cell wall), creating a bright outline in the image^15^. Researchers have developed algorithms to identify the cell boundary by either setting a threshold on the brightness of the membrane-stained cells or by locating the brightest contour^16,17^. To assess the errors and biases in our estimation of cell boundaries from images of membrane-stained cells, we generated digital images of bacteria stained with a membrane marker. When we isolate the effects of projection and diffraction in synthetic images of a membrane-stained cell, we observe that projection causes a notable shift of 3D intensity towards the cell’s centre. This shift is further illustrated in Fig. 2b, in the corresponding radial intensity profile. Such a shift leads to an underestimation of cell dimensions, especially cell width, which is typically estimated from the interpeak distance. Diffraction further exacerbates this intensity shift towards the centre, resulting in an even greater underestimation of width. The magnitude of this shift is influenced by the imaging wavelength, as expected due to diffraction (as shown in Fig. 2c). Note: In this discussion, we have focused on cell width estimation from the interpeak distance of radial intensity profiles of such images, due to its pronounced error sensitivity compared to cell length and its quadratic influence on cell volume, significantly affecting overall size estimation.

Our digital experiments indicate that in a typical widefield imaging setup using a 515 nm fluorescent dye using a 1.45 NA objective, the width of a 1 µm wide cell is underestimated by approximately 20%, and a 0.5 µm wide cell is underestimated by 40% (Fig. 2c). The extent of underestimation is higher for dyes with longer wavelength (Fig. 2c) and for cells narrower than 0.6 µm in width, stained with a red fluorescent dye (emission wavelength = 700 nm), two separate peaks are not observable due to diffraction blur (Fig. 2i), so the width cannot even be estimated. These findings underscore the major biases and limitations of this method for cell size estimation in existing literature.

To validate the predictions from our digital experiments, we labelled the peptidoglycan layer of individual *E. coli* cells with two distinct stains emitting at different wavelengths (HADA = 450 nm and RADA = 580 nm), both being fluorescent D-amino acids (FDAAs). We expect these stains to integrate into the same location in the peptidoglycan layer. However, radial intensity profiles revealed a notable inward shift in the intensity peaks of the longer-wavelength dye (RADA) compared to the shorter-wavelength counterpart (HADA) (inset - Fig. 2d), consistent with our simulated profiles in Fig. 2c. Analysing 137 cells, we found that RADA, with the longer wavelength, led to significantly higher underestimations of cell width compared to HADA (see Fig. 2e, with additional image examples in Supplementary Information 2). These results validate our prediction about diffraction effects on the width estimation. However, since the effects of diffraction on the peak position is dependent on the extent of projection, these effects cannot be eliminated by deconvolution using the PSF, as 2D deconvolution is unable to eliminate the projection effects (shown in Fig. 2f and Supplementary Information 3). 3D deconvolution would partially address this problem, but it is not compatible with single-plane widefield images that are typically acquired during time-resolved imaging of microcolonies. Instead, using superresolution imaging where diffraction effects are minimised (such as SIM, STED, or PALM^18–21^) could help, or employing an imaging system with a shallower depth of field compared to the cell depth (such as a confocal microscope) could reduce the effects of projection and mitigate the resulting shift from diffraction effects (detailed in Supplementary Information 4).

### Errors and biases in size estimation from images of cells uniformed filled with markers

Alternatively, researchers often use thresholding algorithms to segment bacterial cell images based on uniformly distributed fluorescence of molecules within the cytoplasm^22–27^. Various thresholding algorithms are employed to segment cells from their fluorescence images, but each has biases and sensitivities that are challenging to quantify and correct for (Supplementary Information 5). To quantitatively assess the impact of projection and diffraction on extracting cell dimensions from these types of images, we rely on estimating cell width from the full width at half maximum (FWHM) of the radial profile of the intensity.

Unlike in our previous analysis of membrane-stained cells, projection and diffraction have opposite effects on size quantification for cytoplasm-stained cells. Projection effects cause the intensity distribution to shift towards the centre, leading to a bias towards underestimation of cell width from FWHM, while diffraction effects result in light bleeding out, making the radial profile wider than their projected version (see Fig. 2g and h). We demonstrate that increasing the depth of projection leads to an underestimation of cell dimensions beyond a critical cell width (Supplementary Information 4), while higher imaging wavelengths result in increased image blur from diffraction leading to a bias towards overestimated dimensions. The diffraction effect is also apparent in brightfield/phase-contrast images. In Supplementary Information 6, we compare radial profiles of phase-contrast images of the same cell collected with different emission filters. The results reveal that phase-contrast images collected with blue emission filters exhibit significantly sharper images and narrower profiles than those collected with red emission filters. Results from image segmentation of phase-contrast or brightfield images of bacteria are affected by such biases and should be corrected for^28,29^.

It is important to note that both imaging approaches (membrane stained or cytoplasm labelled) exhibit biases in cell dimension estimation that strongly depend on the actual cell dimensions. Fig. 2i shows that the relative width-estimation bias from the membrane image decreases as cell width increases, while the estimates from the cytoplasmic marker exhibit an opposite, but less pronounced effect. In the case of membrane-stained images, the shifts from projection and diffraction happen in the same direction, while they oppose each other in case of cytoplasmically stained images. An accurate model of the imaging system can be used to calculate correction factors for a given wavelength, which could then be applied to estimate the true dimensions from the observed profile. Virtual-microscopy platforms, such as SyMBac, could be utilised to simulate these effects to computationally estimate such correction factors (Supplementary Information 7). However, it is difficult to recover the outline of an individual cell to accurately estimate size and shape using this approach. In the following section, we explore methods for incorporating these effects into training deep learning image segmentation models, enabling the models to accurately estimate cell sizes and shapes from 2D images.

### Deep learning approaches for precise quantification of cell dimensions

There has been a surge in the popularity of deep learning approaches for cell image segmentation^30–35^. However, the accuracy of these models is inherently linked to the quality of their training data. Generating training data for microscopic images of microbes presents unique challenges compared to standard computer vision tasks. Here, projection and diffraction effects are comparable to object dimensions, and as a result impede computational boundary identification, as mentioned in our previous work^12^. Manual annotation is also affected because 2D images lack clear boundaries and contain intensity gradients. In essence, the images are blurry. To evaluate the performance of human annotators, we provided them with synthetic images with accurate ground truths and conducted a benchmarking experiment. Supplementary Information 8 details this experiment and corresponding results, revealing that human annotator performance is not only highly variable, but also consistently exhibits an underestimation bias stemming from projection effects.

Inaccuracies and biases in training data, whether originating from computational thresholding or human annotation, compromise the integrity of object-image relationships, thereby leading to corrupted performance of deep-learning models. The subsequent analysis shows that the highly versatile Omnipose algorithm (specifically bact_fluor_omni)^30^, when trained on human-annotated synthetic fluorescence images, compromises its efficacy in cell segmentation (Fig. 3). This phenomenon parallels findings from our recent publication^12^, where we demonstrated that the segmentation outputs from pretrained models inherit biases from their training datasets, resulting in significant variability in segmentation outcomes and marked deviations from the ground-truth distribution.

**Figure 3-.**
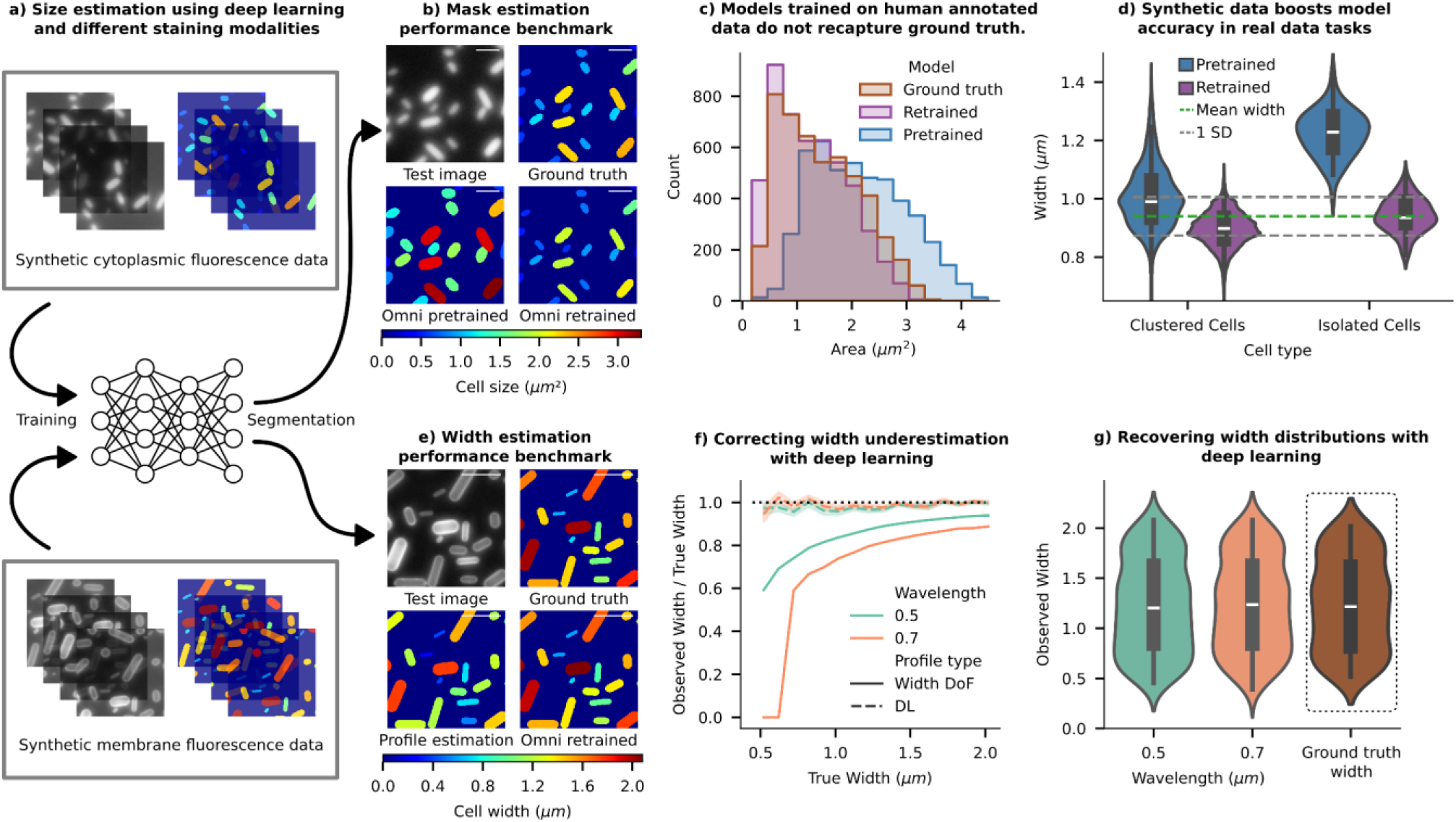
Accurate boundary estimation with deep-learning models trained with synthetic images: **a)** SyMBac can be used to generate synthetic training-data, containing realistic images and accompanying perfect ground-truth, to retrain image segmentation models. Illustrative examples of synthetic image and ground-truth pairs are shown for training Omnipose to learn cell masks from images of cytoplasm-stained samples (top) and images of membrane-stained samples (bottom). **b)** Using synthetic data as a ground truth, we can check the performance of the pretrained bact_fluor_omni model. To alleviate the effects of human annotation quality, we retrained the model on samples of simulated agar-pad data generated using SyMBac. Examples of validation data, with ground-truth masks, and mask outputs from the pretrained and retrained models are shown. To compare between ground-truth masks and output masks, each is coloured based on its total area, and the colormap is given below (scale bar = 2 µm) **c)** Comparison of the output distribution of cell sizes shows that the pretrained model does not reconstruct the underlying ground truth distribution, whereas the output distribution from the SyMBac trained model more closely mimics the underlying distribution. **d)** To show that synthetic data also boosts segmentation accuracy on real data, we analysed patches of densely packed cells to find groups of cells aligned across their long-axis. Since cell width is tightly controlled, we can use these patches of aligned cells to estimate a value for the true population mean width (full analysis is given in Supplementary Information 10). We then generated training data matching the real data’s experimental parameters, and retrained Omnipose. The resulting distribution of widths for isolated cells, and cells within dense colonies is plotted for both the pretrained and re-trained model, showing that re-training on synthetic data makes width estimation more accurate. (Ground truth: 0.94 ± 0.066 μm, pretrained: 1.2 ± 0.10 μm, SyMBac trained: 0.94 ± 0.062 μm). **e)** Using synthetic data of membrane stained cells as a ground truth, we trained an Omnipose model to segment cells. We compared the output widths to those widths measured by calculating the interpeak distance between the labelled cell walls/membranes, as shown in Fig. 2. (Mask colour represents cell width, and the colormap is given below, scale bar = 2 µm). **f)** The fractional underestimation of a membrane stained cell’s width (given by the interpeak distance) is highly dependent upon the width itself, and the imaging wavelength. This is true for a cell imaged in widefield, where the DoF is approximately equal to its width (Width DoF in the legend). Training Omnipose on synthetic data of membrane stained cells makes the deep-learning model (DL) become insensitive to the scale of the cell, as well as the imaging wavelength, unlike the interpeak distance method (error bar = 1 SD). **g)** Comparison of the output mask width distribution of the two simulated datasets to the ground-truth mask width distribution shows that when trained on appropriate synthetic data, the entire population distribution can be faithfully reconstructed irrespective of the imaging wavelength.

The virtual-microscopy pipeline offers an advantage in addressing the issue of user subjectivity and bias in training data. One can generate realistic synthetic microscopic images of microbes accompanied by accurate ground-truth information (Fig. 3a). Training deep-learning models with such synthetic training data enables the models to learn precise object-image relationships (detailed in Supplementary Information 9, Supplementary Information 10, and ref^12^) and mitigates the problem of inaccuracies and user subjectivity in traditional training data. The same Omnipose model, when retrained with synthetic data, produced a segmentation output that more accurately predicted the ground-truth information in the test data, as demonstrated in the ground-truth mask comparisons (Fig. 3b) and input-output size distributions (Fig. 3c). The comparison of cell size distributions indicates that Omnipose training data contains enlarged cell masks.

To experimentally verify and validate the enhanced performance of the retrained Omnipose model compared to the pretrained version, we devised a new assay which leverages the tight width regulation of bacteria^36^. This involved placing a high density of cells on agar pads, capturing images of both isolated cells and cell clusters, and then estimating the ground-truth widths of individual cells based on their average width in aligned space-filling patches (further explained in Supplementary Information 10). The average cell widths estimated from the patches were tightly distributed (0.94 ± 0.066 μm). The estimated mean width from the patch analysis should match the average widths of isolated individuals, as the cells were not grown on the agarose pad, and therefore not allowed to differentially adjust to the imaging environment. Subsequently, the widths obtained from this analysis were compared with those derived from the segmentation outputs of both the pretrained omnipose model (1.2 ± 0.10 μm) and the retrained model using synthetic images (0.94 ± 0.062 μm). The results demonstrate that the retrained Omnipose model exhibits both higher precision and accuracy in estimating cell widths when compared to its pretrained counterpart (Fig. 3d). The comparison of masks presented in Supplementary Information 11 reveals that the original Omnipose model generates substantially larger masks for isolated cells than for cells within clusters, resulting in significant variability and bias in the predicted cell width. In contrast, the output masks from the Omnipose model retrained with synthetic data demonstrate robust performance.

Motivated by these results, we explored an additional application of this approach in the analysis of membrane stained images. We retrained the Omnipose model with pairs of synthetic fluorescent images of membrane-stained cells and corresponding ground truths (Fig. 3a - bottom). The estimated cell outline from the contour of the membrane stained images (as described in ref^37^) significantly underestimated the cell area compared to the ground truth masks (Fig. 3e). The relative error in width estimation was size and wavelength dependent (Fig. 3f), consistent with the previous discussion. Conversely, the comparison of output masks from the retrained Omnipose and the ground-truth cell mask illustrates high accuracy and precision of the deep-learning model. The model robustly learns the offset created by diffraction and projection as a function of size, and the estimated width closely tracks the ground truth across a wide range of input widths and in a wavelength independent manner (Fig. 3f and g). The combination of these digital experiments and real experiments illustrates how synthetic training images can capture the subtle effects of projection and diffraction and augment our capabilities of estimating true cell sizes using deep-learning models.

### Quantifying Fluorescence Intensities of Individual Cells

Next, we address the issue of quantifying the intensity of individual cells from their fluorescence images. Measuring the total fluorescence intensity of labelled molecules within a cell is crucial for estimating their abundance. This capability enables researchers to monitor the dynamics of cellular processes using time-resolved single-cell image data^38^. The variation in signal intensities among individual cells within the population and over time offers insights into the key regulatory variables and noise sources^39^.

Usually, for the sake of experimental simplicity, microcolonies of microbial cells are cultivated on agarose pads^27^. This setup enables the tracking of individual cell intensities over time and comparison of intensities among colony cells at different time points. However, such experimental designs, including microfluidic devices with densely packed cells^40,41^, introduce a significant artefact in single-cell intensity measurements due to a combination of diffraction and projection effects from the imaging system. The PSF of an imaging system disperses light away from its source. In the context of a cell filled with fluorescent emitters, the emitted light extends beyond the true cell boundaries, making solitary cells appear dimmer (see Fig 4a and 4b). In densely packed clusters, the dispersed light is erroneously attributed to neighbouring cells. We previously termed this phenomenon as ‘light bleedthrough’^42^. Light bleedthrough substantially distorts intensity estimates of cells within a colony, leading to misinterpretations of the strength and noise in gene expression levels, as explained below.

**Figure 4-.**
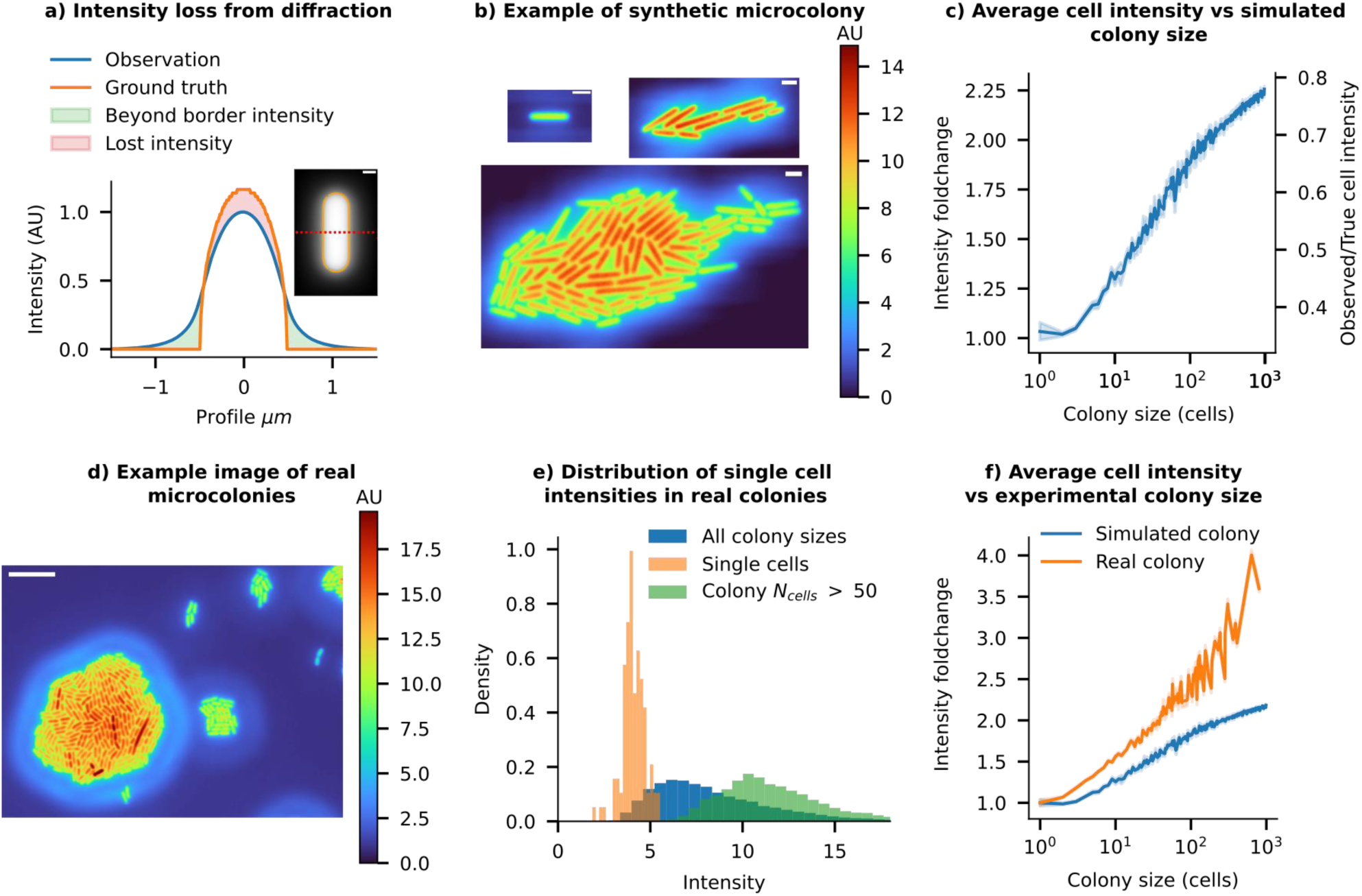
Colony size affects single cell intensity quantification - real and synthetic data: **a)** The radial intensity profile of a simulated fluorescence cell illustrates how intensity is lost from the cell contours due to diffraction effects. (Scale bar = 0.5 μm) **b)** Example snapshots of simulated fluorescence images of a growing microcolony using a 100x 1.49NA objective’s instrumental point spread function (iPSF). All cells in the simulation have identical ground truth intensities. Isolated cells and individual cells in small microcolony sizes show low cell intensities, while as the colony grows larger, cells in dense regions, such as at the centre of the colony, begin receiving more light contribution from surrounding cells, artificially increasing their perceived intensity. (Scale bar = 2 μm) **c)** As colony size increases, the mean observed intensity of each cell in the colony also increases (Error bars = 99% CI). Relative changes in intensity compared to isolated individual cells are shown on the left-hand y-axis. Cells approach the true mean intensity as the colony size increases, as shown on the right-hand y-axis. **d)** A false-coloured image of cells on a real agarose pad showing ‘preformed’ microcolonies of various sizes, along with a single cell (white arrow). The relative intensity scale is shown on the right. Important features to note are the similarity to the simulated data shown in b) (Scale bar = 10 μm). **e)** The intensity distribution of cells depends on the size of clusters they belong to. Isolated individual cells have low mean intensities, while cells from preformed microcolonies with more than 50 cells have 3x higher mean intensities. **f)** Experimental data shows that the average intensity of cells increases with the population size of microcolonies (shown in orange). The trend from simulated colonies is shown in blue.

With the SyMBac virtual microscopy system, we can quantify these effects and verify them through experiments on real microcolonies (see example images of synthetic microcolonies in Fig. 4 real examples in Fig. 4d). Crucially, while measuring the instrumental PSF (iPSF) of one’s microscope is a standard procedure, they are not typically imaged over a domain large enough to capture the effects of light bleedthrough at long distances (> ∼15 μm for high NA objectives), since the signal to noise ratio becomes low. Thus, we pursued analytical fits to the iPSF to extend its range for simulating long-range diffraction effects. The most suitable method involves extracting the pupil, followed by reconstructing a phase-retrieved PSF which includes appropriate aberrations^43^. However, as detailed in Supplementary Information 12, while the reconstructed PSF effectively captured the aberrations in our system, it failed to replicate the long-range effects observed in the iPSF. A theoretical PSF (tPSF) model^44^ gave a much better fit to the entire iPSF (Supplementary Information 13). However, since we are interested in simulating the long range effects of diffraction, one must extrapolate the function domain of the fitted PSF. We found that the tPSF did not extrapolate well when fitted to a crop of the iPSF. We therefore resorted to an ad hoc empirical function fit, which we call an “effective” PSF (ePSF), which we verified was able to extrapolate to the entire function domain despite being fitted on only a small crop of the iPSF (see Supplementary Information 14). All simulations of light bleedthrough effects in microcolonies were carried out using this ePSF model.

### Colony size affects single cell intensity quantification

Our simulations suggest that due to the loss from light bleedthrough, an isolated cell appears only 30% as bright as its true intensity (Fig. 4c). 70% of the intensity is lost to the surroundings, and can end up in nearby cells. As the microcolony size increases, more neighbouring cells contribute to the intensity of cells within the colony through the light bleedthrough effect. Consequently, the mean intensity of cells within a simulated microcolony rises monotonically with colony size, reaching 70% of the true intensity in very large colonies (>1,000 cells, see Fig. 4b and c). As the colony size tends to infinity, the mean intensity of an individual cell should converge to the true mean intensity, as all the lost intensities are allocated to other cells within the colony. These simulations predict that individuals within a colony of a hundred cells should appear, on average, 2-2.5 times brighter than isolated cells (Fig. 4c).

To validate these predictions, we conducted experiments with microcolonies on agar pads, comparing the intensity of cells within colonies of different sizes with that of isolated cells. To ensure a consistent intensity distribution among all cells, we placed a high density of cells on agar pads and captured instantaneous images of cell clusters, as opposed to allowing colonies to form gradually, which could introduce temporal intensity variations. This method allowed us to obtain ‘preformed microcolonies’ of varying sizes while maintaining the original cellular content distribution. As expected from the analysis of simulated microcolonies, snapshots of real cell clusters clearly show higher intensity in cells from larger colonies compared to isolated cells (see Fig. 4d). The intensity distributions of cells in large ‘preformed colonies’ (number of cells >300) do not overlap with those of isolated individuals (see Fig. 4e). The trend in Fig. 4f, which illustrates the mean intensity of cells relative to their preformed colony sizes, is qualitatively similar to the trend predicted from the digital experiments. Nevertheless, the magnitude of bleedthrough effects witnessed in the experimental microcolonies exceeds that of the simulated colonies. This disparity may arise from the mismatch between the long-distance tails of the ePSF and the actual iPSF of the system, the possible contribution of scattering in the imaging media, or field-dependent aberrations not captured in our simulations.

### Light bleedthrough affects noise and correlations in single-cell intensity measurements

The light bleedthrough effect goes beyond its impact on mean intensity, introducing subtle local variations in individual cell intensities. Since the degree of bleedthrough depends on the number of neighbouring cells, the intensity of individual cells varies based on their position within the colony (Fig. 5). Cells closer to the centre, receiving contributions from more neighbours, appear brighter in images than those near the edges (see Fig. 4b and d). Consistent with the predictions from our digital simulations of microcolonies, the experimental data reveal a correlation between spatial intensity patterns, the number of neighbouring cells, and intra-colony position, (see Fig. 5a-c). Additional example images of microcolonies from imaging experiments are shown in Supplementary Information 15.

**Figure 5-.**
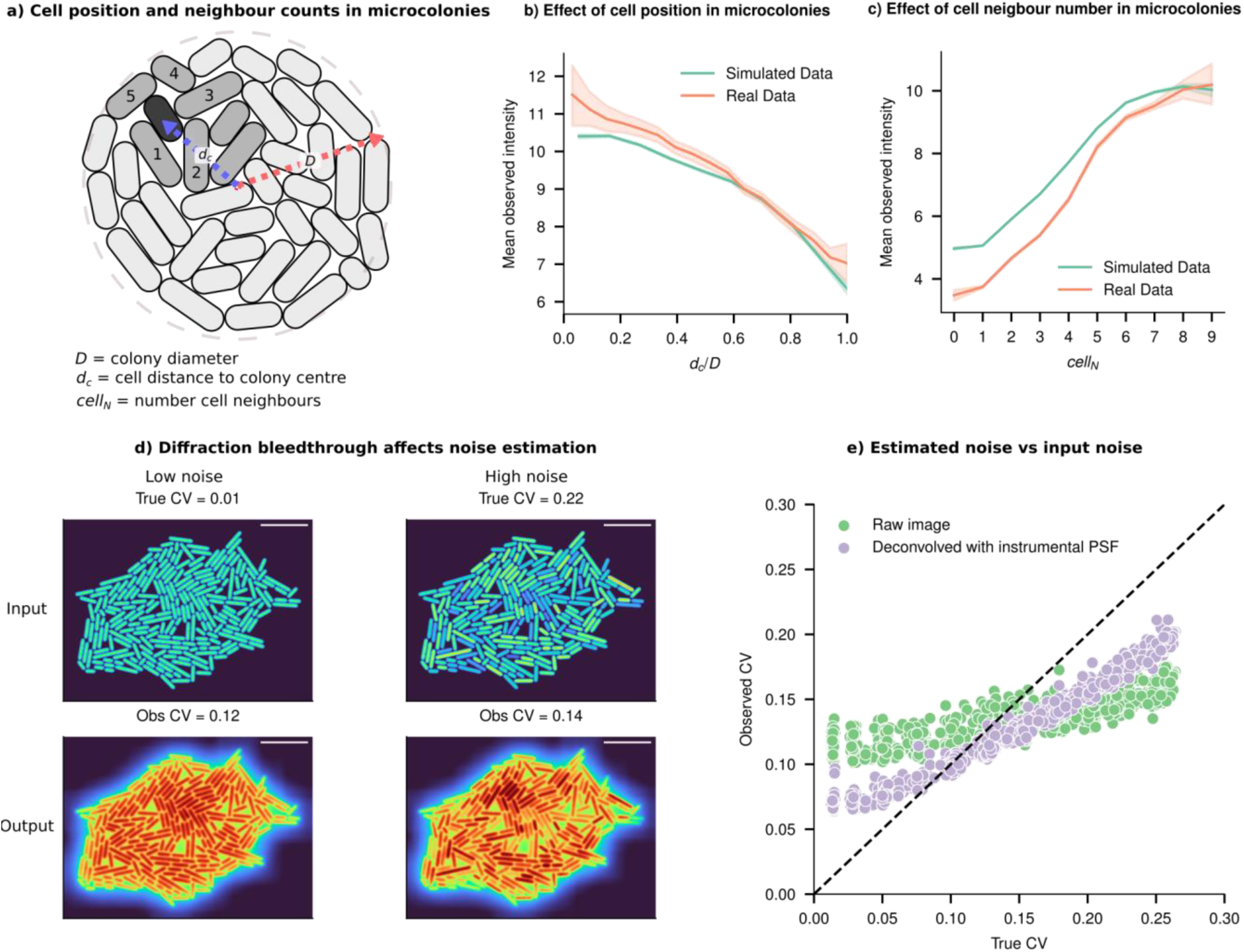
Light bleedthrough corrupts estimates of correlation and variance in single cell intensities: **a)** Schematic representation of a cell (green) some distance from the centre of the microcolony, with its neighbouring cells labelled (lighter green). The intensity of individual cells depends on their position within the microcolony, given by *d_c_/D*, where dc is the distance from the colony centre, and *D* is the colony diameter and the number of direct neighbours (*cell_N_)*. **b)** Cells closer to the centre of a simulated colony appear brighter than cells at the periphery due to light bleedthrough (Error bars = 99% CI, data sample averaged for colonies of size 20-1000 cells). The position-dependent trend predicted from simulated microcolonies (green) is consistent with experimental results (orange). **c)** Simulated cells with more neighbours appear brighter (Error bars = 99% CI, data sample averaged for colonies of size 20-250) consistent with experiments (orange). **d)** Two simulated colonies with CVs of 0.01 and 0.22, respectively, along with their convolution with the iPSF, showing that their observed CVs are very similar despite markedly different underlying cell intensity distributions. e) At low noise (true CV < 0.15), where the underlying cell intensity distribution is uniform, the PSF causes artificial position-dependent variation in cell intensities (increasing CV). Conversely, when the input CV is high (true CV > 0.15), the PSF acts as a blurring filter, lowering the variance in the population by allocating intensities from brighter cells to neighbouring dimmer cells (lowering CV).

Such phenomena, where intensity of individuals appear to be dependent on their position or number of neighbouring cells, can lead to misinterpretations in quantifying intensity correlations and cellular heterogeneity^5,8^. It may wrongly suggest that an individual cell’s intensity is influenced by interactions with neighbouring cells, incorrectly implying non-existent biological mechanisms. In studies using intensity distribution patterns as evidence of cell-cell interactions, researchers should consider the confounding influence of optical effects and implement appropriate controls to differentiate genuine biological interactions from optical artefacts^7^. The use of digital control experiments via virtual microscopy platforms, like SyMBac, can help identify potential artefacts specific to a given experimental design, including optical specifications and sample configurations.

Light bleedthrough effects also cause a major artefact in noise estimation from single-cell data, which is somewhat counterintuitive. In the absence of true population variability (coefficient of variation, CV = 0), positional factors within the microcolony and the number of neighbouring cells can artificially introduce variability and result in a higher estimated CV (Fig. 5d). Conversely, when substantial variation exists among cells, the PSF acts as a smoothing filter, redistributing intensity from brighter to dimmer cells (see Fig. 5d and e), leading to an underestimation of inherent variability. These paradoxes emphasise the complexities introduced by diffraction effects in the temporal quantification of gene-expression variability. They produce similar results across different underlying distributions (further examples in Supplementary Information 16) and present challenges for correction because the effect’s magnitude and direction depend on unknown ground truth CV, as well as the size and shape of the microcolonies.

It is conceivable that one could leverage deconvolution to correctly assign light to specific pixels in the image. In actual image formation, the Point Spread Function (PSF) has an infinite range, leading to long-range diffraction effects accumulating within dense microcolonies. This results in diffracted light potentially ending up several pixels away from the original point source. However, deconvolution methods applied in the literature sometimes use kernel sizes far smaller than the data, and thus merely results in a sharpening of the image, failing to accurately reassign light from beyond the kernel’s boundaries. This phenomenon is illustrated using experimental and simulated images in Supplementary Information 17 and Supplementary Information 18. As shown in Fig. 5e, where deconvolution is performed with a kernel measuring 125×125 pixels, only marginal improvement in noise estimation accuracy is seen. Ideally, the deconvolution should use a kernel size as large as the data being deconvolved. Since such a large iPSF is unattainable, deconvolution with the full ePSF was performed. While it gave a marked improvement over the iPSF, it was unable to fully recover the underlying ground truth intensity distribution in experimental data (Supplementary Information 17).

### Microfluidic imaging platforms for robust intensity quantification

Given the challenges associated with accurately knowing the PSF and exact configuration of cells within a microcolony, the task of estimating the true intensities of individual cells, especially for quantifying noise or correlations, becomes nearly impossible. Additionally, the influence of projection and scattering effects and potential inhomogeneities in the growth environment^45^ is hard to eliminate. Therefore, we suggest that, considering knowledge of diffraction effects, researchers could design their experiments differently. For instance, utilising a structured imaging platform, where cells are maintained at a fixed distance from each other, can help minimise the bleedthrough effects.

To systematically analyse the constraint on such an imaging platform, in our simulations on an array of digital cells, we explored how the extent of intensity bleedthrough depends on the inter-cell distance in such an array (Fig. 6a). The percentage bleedthrough contribution from neighbouring cells is plotted as a function of distance along the short and long axis of cells (x and y, respectively) (Fig. 6b). The heatmap illustrates that in a closely packed array, shown in the left-top corner, the intensity of individual cells receives an additional ∼100% contribution from neighbours, causing cells to appear 2x brighter than isolated cells—a finding consistent with our earlier discussion. To reduce the light bleedthrough effects to <1% of the true intensity, cells need to be at least 10 μm apart from each other (a conservative estimate based on the ePSF).

**Figure 6:**
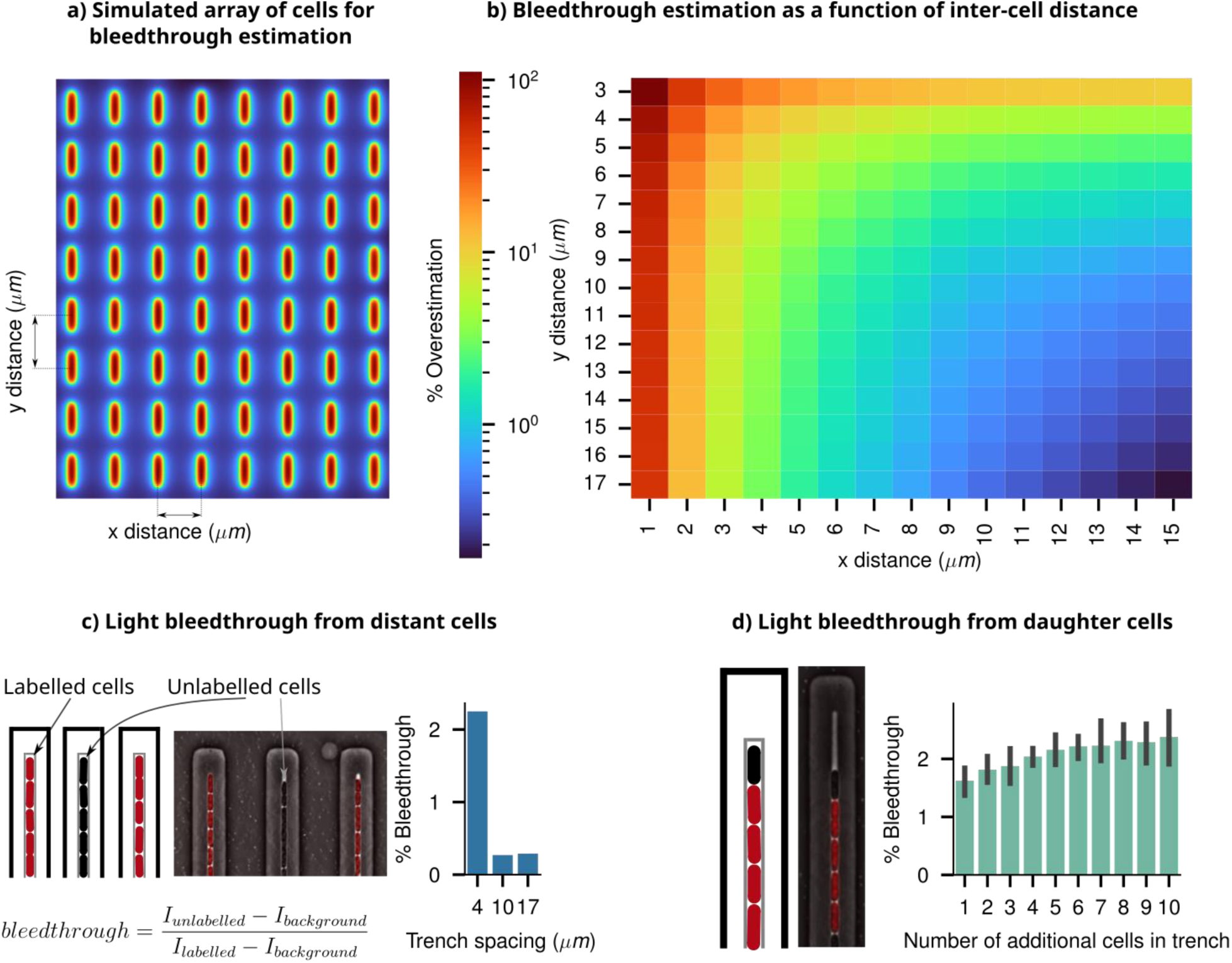
Intensity bleedthrough characterisation using simulated array and microfluidic experiments: **a)** A simulated array of cells with controllable x and y cell pitch. The grid corresponds to the heatmap in panel b, with increased x and y spacing between cells lowering the intensity bleedthrough from neighbours. **b)** The heatmap shows the overall intensity bleedthrough percentage of a cell within an ordered array of neighbours as a function of inter-cell distance. **c)** Characterisation of intensity bleedthrough from distant cells within the mother machine microfluidic device. A schematic representation of the mother machine is given, along with a phase-contrast image taken of the same device, with the (mCherry) fluorescence channel overlaid to indicate labelled and unlabelled cells. Varying the spacing between mother machine trenches affects the amount of light bleedthrough incurred. **d)** Bleedthrough from within a single trench in the mother machine; an unlabelled cell’s intensity is measured and its apparent intensity increases as the number of additional labelled cells within a trench increases, despite it being unlabelled.

Microfluidic devices, such as the ‘mother machine’^2^, provide a viable solution for maintaining cells at a fixed, constant distance from one another. This device keeps cells in short vertical arrays placed at a specific distance apart. By selecting the appropriate spacing between these arrays through specific device design and optimisation, researchers can effectively eliminate the bleedthrough effect, allowing for accurate estimation of the heterogeneity and fluctuation dynamics of intensities from individual cells^46^.

To assess the effect of bleedthrough in these scenarios, we conducted experiments by mixing unlabeled cells with fluorescently labelled cells in the mother machine (black and red coloured cells in Fig. 6c and d respectively). Details of the experimental design, analysis, and results are shown in Supplementary Information 19. Intensities of unlabeled cells, as a function of the number and distance of their neighbouring fluorescent cells, were calculated to estimate the percentage bleedthrough effect. The results from this analysis, along with data from a previous paper using a similar approach^46^, show a quantitative match with the simulated trends. A distance of >10 μm between trenches is sufficient to reduce the extent of bleedthrough to below ∼1%. Microfluidic device design and performance verification using digital microscopy experiments should be routinely employed to estimate and eliminate unwanted optical effects in microscopy data.

### Counting Single Molecules

Quantifying the abundance of low-copy tagged molecules introduces unique challenges. When the collective fluorescent signal from these tagged molecules approaches the background autofluorescence of the cell, interpreting intensity values in terms of abundance becomes complex. In the case of species with moderately low copy numbers per cell (approximately 50-100 copies per cell), some researchers have employed techniques like background deconvolution^4^. However, these methods fall short of achieving single-cell resolution since they deconvolve the entire distribution along with autofluorescence levels. Moreover, these distribution deconvolution techniques are ineffective for proteins with very low copy numbers (less than 20), which often play critical roles in gene expression regulation, including gene copy numbers, transcription factors (TFs), mRNAs, and plasmids^47^. To reliably quantify the abundance of low-copy proteins, it is essential to count individual fluorescently tagged molecules within a cell^48^.

Accurately determining the copy number of labelled molecules remains a formidable task due to the interaction of the diffraction limit and the 2D projection of 3D-distributed emitters. In 2D images, individually tagged molecules manifest as diffraction-limited, blurry spots, with the extent of blur contingent upon their distance from the focal plane. Consequently, when spots are positioned closer to each other than the resolution limit, (in the XY plane, regardless of their position in Z, due to projection), they can merge into a single spot in the projected image. Adding to the complexity, the 3D characteristics of the PSF make it difficult to detect out-of-focus spots (Fig. 7a). Both of these effects collectively contribute to an underestimation of the molecule count, even when there are only two copies per cell.

**Figure 7:**
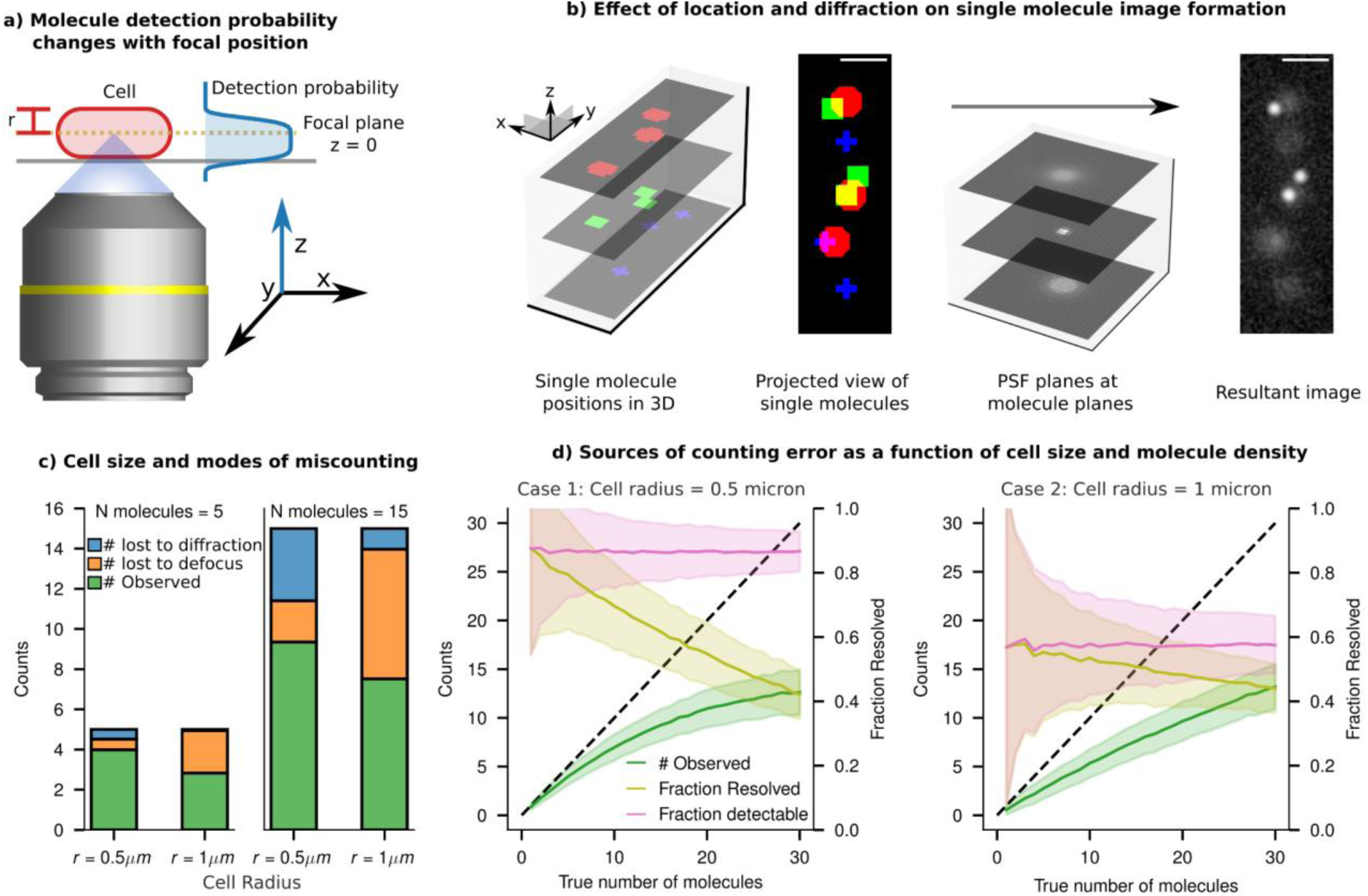
single particle counting errors from the combined effect of projection and diffraction: **a)** Schematic of an epifluorescence single molecule imaging setup, where the position of emitters within the cell determines the extent of defocus in their image and determines their detection probability. **b)** Molecules in a cell exist in three dimensions. Shown are 9 molecules in a digital cell and their ground truth positions projected in xy. Upon convolution with the PSF, the resultant image is a projection of all 9 molecules, of which only 3 are in focus. The remaining 6 molecules are dim and hard to detect and two of them are very close. **c)** Simulated sampling of points within a digital cell shows ‘naive’ detection bounds and sources of undercounting. If points had a SNR lower than the 99th percentile of the background PSNR, they were considered too dim to detect. Additionally, if two molecules were within 1 Rayleigh criterion of one another, they are considered too close to resolve, and thus these molecules are lost due to diffraction. The remaining population of molecules are considered countable. **d)** Relative contributions of each mode of undercounting (lost to diffraction and lost to defocus) are plotted as a function of true count of molecules in a narrow cell (r = 0.5 μm) and a thicker cell (r = 1 μm). The resolvable fraction decreases rapidly with increasing density of molecules, whereas the detectable fraction stays constant.

In Fig. 7b, we illustrate the extent of undercounting from various sources using digital imaging experiments. We employ a “naive counting” criterion, as described in the Methods section, which includes the enumeration of molecules that are out of focus, those that are undercounted due to proximity to diffraction-limited spots, and the cumulative count of both individual and cluster molecules perceived as singular due to the diffraction limit. This approach allows us to identify error sources that are influenced by cell dimensions (spot density and projection) and experimental setups (diffraction and depth of field). The results in Fig. 7c show that when the count is small (n = 5), most spots are isolated, resulting in minimal losses due to diffraction effects. However, increasing the cell thickness (from cell radius = 0.5 μm to radius = 1 μm) leads to a significant fraction of spots becoming undetectable as they get blurred and dimmed due to the defocused PSF. On the other hand, as the molecular count increases, there is a corresponding increase in the proportion of molecules that are undercounted due to the diffraction limit, since an increase in spot density leads to increase in the fraction of spots being unresolvable from their neighbours. However, the proportion of molecules lost due to defocus remains constant, dependent solely on the volume fraction of the cell situated outside the focal plane (Fig. 7d).

This digital experiment reveals that the combined effects of projection and diffraction lead to substantial undercounting, even for molecules present in very low quantities: in an average-sized bacterium, a single snapshot may incorrectly count the only two molecules as a single entity approximately 5% of the time (Supplementary Information 20). The proportion of undercounting escalates rapidly with the increase in number of molecules present per cell, as depicted in Fig. 7d, and for a copy number of 15 molecules/cell, one underestimates by 40%.

### Smart-microscopy approaches for improving counting performance

The term ‘smart-microscopy approaches’ denotes utilising domain knowledge of a specific imaging system and subject to craft targeted microscopy solutions, encompassing both acquisition and analysis. To improve counting performance, the knowledge of the depth-dependent detection probability and the cell volume can be leveraged to calculate a correction factor, addressing the loss of molecules due to defocus. At the focal plane, a molecule will be most in focus, and given that it is bright enough, will exceed a threshold SNR for detection. The probability to exceed this SNR threshold decreases as the molecule shifts out of the focal plane. We call this probability function *D*(*z*). We derived this empirical depth-dependent detection probability function for our imaging system from the instrumental PSF (shown in Fig. 8a and detailed in Supplementary Information 21).

**Figure 8:**
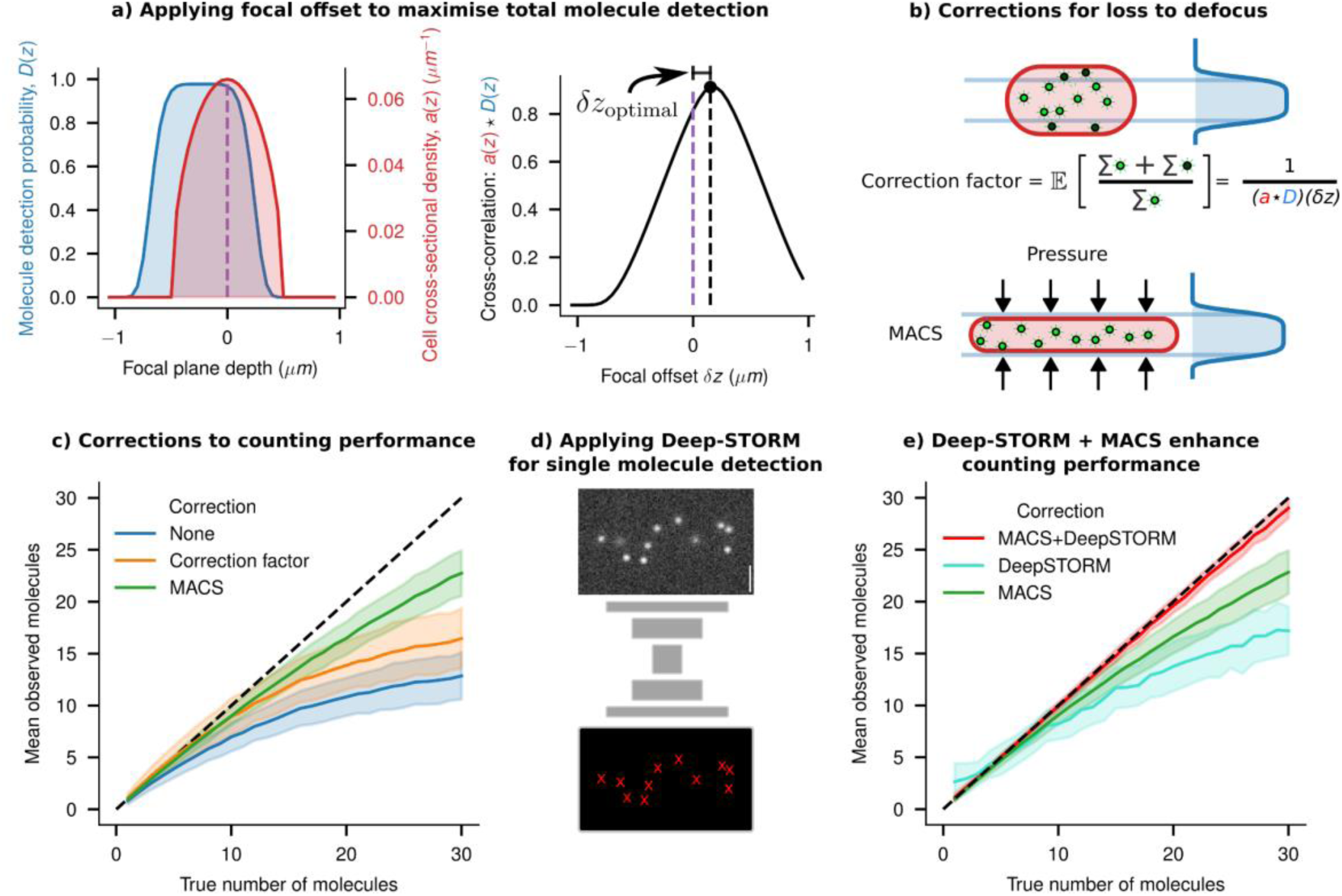
Smart microscopy approaches to improve counting performance: **a)** The depth dependent probability function *D*(*z*) of an imaging system is shown in blue. The cell’s cross-sectional density is given in red as *a*(*z*). The overlapping area between these two profiles (given by the cross-correlation *a*(*z*)\**D*(*z*) gives an estimate of the fraction of the molecules which will be observed. To maximise the number of molecules detected, one can shift the objective by an optimal amount, given by *δz*_optimal_, which is where the two functions have maximum overlap. **b)** A schematic showing a cell in the optimal focal position relative to the detection probability function, thus detecting the maximum number of molecules possible. The true number of molecules can then be estimated by multiplying the observed count (green) by a correction factor (accounting for the lost molecules, shown in black), which intuitively is the reciprocal of the overlapping area (full derivation in Supplementary Information 21). Another approach to detecting more molecules is to modify *a*(*z*) by physically compressing the cell (using Microfluidics-Assisted Cell Screening (MACS)) bringing the entire cell’s volume within the maximum region of *D*(*z*). **c)** Applying the correction factor or compressing the cell using MACS improves the counting performance compared to the naive estimate. Both of these approaches reduce the error from defocus, but undercounting errors at higher counts happen due to diffraction effects. **d)** A schematic architecture of the Deep-STORM single molecule localisation network is shown, which was trained using synthetic single molecule images. **e)** Applying Deep-STORM to molecule counting improves performance, but combining it with MACS leads to near-perfect detection and counting up to a higher density of molecules.

Upon closer examination, we observed that this function is offset from the objective’s focal plane because of the asymmetric nature of the iPSF along the Z axis. To maximise the number of detectable molecules within a cell, it is necessary to optimise the overlap between the cell’s cross-section and this function. A cell’s cross-sectional density is given in red as *a*(*z*) (Fig. 8a). The integral of *a*(*z*) between two z positions within the cell will give the volume fraction, and hence the fraction of molecules between those positions (assuming a uniform distribution of emitters within the cell). The detection probability *D*(*z*) can be shifted by focussing the microscope’s objective up and down. Thus, the overlapping area between these two profiles (given by the cross-correlation *a*(*z*)\**D*(*z*) gives an estimate of the fraction of the molecules which will be observed, for all shifts of *D*(*z*). To maximise the number of molecules detected, one can shift the objective by an optimal amount, given by *δz*_optimal_, which is where the two functions have maximum overlap (Fig. 8a).

Once the focal plane is adjusted, we can compute the fraction of molecules lost to the out-of-focus volume to find an empirical correction factor (Fig. 8b). This correction leads to improved counting performance (but only on averaged counts), as demonstrated in Fig. 8d. Alternatively, using microfluidic platforms to flatten cells can bring a larger number of spots into focus (Fig. 8b). Additionally, the expanded cross-section of the flattened cells in the imaging plane slightly reduces the undercounting effect caused by the diffraction limit (Fig. 8c), consistent with previous findings in the work of Okumus et al. (Microfluidics-Assisted Cell Screening - MACS)^10,49^.

To further improve the counting performance, we explored the potential of designing an image analysis pipeline that leverages knowledge of defocus and spatial patterns from simulated data to enhance counting accuracy. To pursue this, we retrained Deep-STORM, a well-established deep-learning network designed for super-resolving single molecule images^50^. Deep-STORM leverages a convolutional neural network architecture to super-resolve single molecules based on local intensity patterns and spatial relationships. We have trained the Deep-STORM model using simulated synthetic images, which contains a varying number of spots with appropriate defocus depending on their position within the digital phantom cells (Fig. 8 d). This training enabled the model to consistently and accurately count molecules to a larger copy-number (Fig. 8e) compared to the naive counting estimates and the performance demonstrated in previous research in this field^10^.

Just as observed with deep-learning models used for cell segmentation (discussed in the “Deep-learning approaches for informed cell segmentation” section), training these models with realistic synthetic data significantly enhances their ability to detect single molecules. Our previous analysis demonstrates that flattening the cells using platforms like MACS reduces the fraction of defocused spots and emulates a situation where most spots are in focus (Fig. 8b). Indeed, Deep-STORM models, when implemented in a simulated MACS type scenario, performed reliably up to a very large number of molecules per cell (> ∼25 molecules/cell, Fig. 8e).

## Discussion

The advancement of quantitative microbiology relies heavily on the accurate interpretation of microscopy data. We have employed virtual microscopy experiments and targeted real experiments to systematically explore the challenges and potential pitfalls associated with using microscopy data to quantify the size and content of microbial cells. Our focus was on projection and diffraction effects, particularly significant for microbial cells due to their size.

Our findings reveal significant impacts of projection and diffraction on the performance of image segmentation algorithms in accurately identifying cell outlines from fluorescence and brightfield images of bacteria. Both traditional segmentation techniques and machine-learning approaches experience biases in cell size estimation. The extent and direction of this bias depends on various factors, including labelling methodologies, imaging configurations, and the cell’s dimensions, which makes it difficult to correct for. However, we found that the bias and error can be mitigated when using machine-learning methods trained with synthetic data that incorporates these effects.

Timelapse imaging methodologies, commonly employing agar pads and microfluidic devices, are frequently utilised for investigating live-cell gene expression dynamics and heterogeneity^39,41,51,52^. Using digital image simulations and experimental fluorescence imaging of cell clusters, we found that the accurate quantification of true cellular fluorescence signals in clustered configurations (’microcolonies’), is difficult due to diffraction-induced misallocation of light intensity from adjacent cells. Such distortions impact both the estimation of expression variation and correlation analyses conducted on these platforms^5,7,8,53^. Deconvolution can improve, but not entirely eliminate, these artefacts, and its fidelity to ground truth strictly depends on the precision and size of the deconvolution kernel. In this case, experimental design changs, such as the use of imaging platforms like microfluidic devices, where cells can be kept at specified distances, can reduce such distortions^42,46^.

Similar challenges arise in the quantification of low copy number moieties, such as mRNA, plasmids, or proteins, complicating the accurate counting of more than five individual molecules. Caution is warranted when interpreting ‘single-molecule’ images and results from estimated molecular ‘counts.’ To address these challenges, alternative experimental designs and deep-learning-based analysis protocols were proposed and substantial improvements in counting accuracy were demonstrated.

In summary, the analysis presented here underscores the critical importance of understanding the artefacts and aberrations incorporated into microscopy data to extract meaningful information about microbiology, whether it involves the shape and size of cells or their content from intensity measurements or single-molecule counting. We advocate for the routine use of digital experiments with virtual microscopy platforms to test limitations of experimental design and potential optical illusions, ensuring ‘informed’ interpretations of imaging data. This knowledge can further inform the design of ‘smart microscopy’ experiments, leveraging domain knowledge to create appropriate imaging platforms and machine-learning models trained with relevant ‘synthetic images.’ The analysis and discussion presented in this paper should guide improved experiment design and help with quantitative interpretation of microscopy experiments in microbiology.

## Methods

### Computational Methods

#### Virtual fluorescence microscopy using SyMBac

In this study, image simulations were conducted utilising the SyMBac Python library^12^. Unless specified otherwise, all virtual fluorescence microscopy images were generated following a consistent workflow: 1. The 3D spherocylindrical hull of a digital cell was positioned within a defined environment—either a) isolated, b) among scattered cells, or c) within a microcolony if colony growth simulation data were available. 2. Fluorescent emitters were uniformly sampled within the cell volume and indexed within a 3D array, the value at each index denoting the emitter count. For cells with homogeneously distributed fluorescence, a “density” value was established, defined as the average number of emitters per volumetric element within a cell. Thus, the total number of molecules within a cell was calculated as the product of this density and the cell volume. 3. Subsequently, diffraction and projection effects were simulated through the convolution of this dataset with a point spread function (PSF). Convolution was executed employing either a theoretical, effective, or an instrumentally measured PSF (tPSF, ePSF, or iPSF respectively). The PSF_generator class within SyMBac was used to generate synthetic PSFs in accordance with the model from Aguet^44^. To simulate the projection and out-of-focus light characteristic of a widefield fluorescence microscope, the centre of the point spread function (PSF) is assumed to be aligned with the midplane of the cell. Each slice of the PSF is accordingly convolved with the corresponding slice of the cell, as illustrated in Fig. 1b, Supplementary Information 22, and Supplementary Information 23. If using a tPSF or ePSF, convolution is done at a high resolution, and then downsampled to the pixel size of the simulated camera in order to capture the high frequency features of the kernel.

To artificially modulate the depth of field of the microscope and thereby mitigate projection artefacts, the number of PSF planes convolved with the cell can be truncated. For instance, in a 1 µm wide cell simulated at a pixel size of 0.0038 µm, there would be 263 slices in the Z-direction. To generate an image devoid of projection artefacts, only the middle Z-slice of the cell is convolved with the middle Z-slice of the 3D PSF. This method is applicable for both simulated and empirically measured PSFs. To eliminate diffraction effects, the PSF can be substituted with an identity kernel, which, upon convolution, reproduces the original data. To modulate the effect of diffraction continuously, the frequency of light employed to simulate the kernel can be arbitrarily adjusted.

#### Simulations of fluorescence images of individual cells

To preclude errors stemming from length underestimation in width assessments, digital cells with a fixed length of 6 μm were utilised. The cell width was manipulated to range between 0.5 and 3.0 μm, while the simulated depth of field was adjusted between 3 μm (for the widest cell) and 0.0038 μm (for a single Z-slice). The PSF generator was configured to “3d fluo” mode for 3D convolution, employing the model from Aguet^44^. Additionally, an identity kernel served as the PSF for a theoretical undiffracted microscope with an imaging wavelength of 0 μm.

To simulate images of cells with cytoplasmic fluorescent markers, 3D spherocylindrical cells were rendered, and emitters sampled as described above. To simulate membrane fluorescent cells, fluorescent emitters were sampled only within a single pixel layer corresponding to the outermost cell volume. We ensured to render images at a high resolution, allowing accurate drawing of the cell membrane. Rendering at a lower resolution, even a single pixel would be significantly thicker than the true thickness of the cell membrane.

Simulated images of fluorescent single cells, either with cytoplasmic markers or membrane markers, were generated at different wavelengths, widths, and depths of focus. It is important to note here that when the depth of field is changed, this is a simulation of a non–physical effect. In actuality, the volume of out-of-focus light captured by the microscope is determined by the objective lens. By adjusting the number of Z-PSF layers with which the 3D cell volume is convolved, the non-physical manipulation of out-of-focus light collection by the microscope’s objective is simulated. Despite its non-physical nature, this is a valuable exercise for identifying sources of measurement bias and error. A similar argument applies to diffraction: the use of an identity kernel simulates an image devoid of diffraction effects. Though non-physical, this is instrumental in examining how an image is compromised by the microscope’s PSF. In real experiments, both projection and diffraction effects co-occur; hence, the comparative analysis is limited to simulated images that incorporate both phenomena.

#### Cell size quantification and analysis

Following the simulation of individual cells, errors in size estimation attributable to diffraction and projection were quantified using two methodologies. For cells marked with cytoplasmic fluorescence, a binary mask of the resultant synthetic image was generated employing Otsu’s thresholding algorithm^54^. Dimensions along the two principal axes of the binary object were then calculated to ascertain length and width. In contrast, for synthetic images featuring fluorescent membrane markers, dimensions were determined by measuring the inter-peak distance along the one-dimensional intensity profile, which was aligned with the two principal axes of the cell.

#### Simulations of fluorescence images of microcolonies of cells

The agar pad simulation feature of SyMBac was used to generate microcolony simulations^12^, each of which was terminated when reaching a size of 1000 cells. All microcolonies are restricted to monolayers. This pipeline leverages the CellModeller platform, which is integrated into SyMBac, for the creation of ground-truth microcolonies^55^. For these simulations, a constant cell width of 1 μm was maintained, with cellular division programmed to occur at a target length of 3.5 μm ± *Uniform*(−0.25μm, 0.25μm) resulting in sequences of densely packed, proliferating colonies.

Individual cells in each colony simulation contain uniformly distributed fluorescent emitters. To control the coefficient of variation (CV) of intensity among the cells within a microcolony, we fixed the mean density of emitters within a cell, but varied the variance of a truncated (at 0) normal distribution in order to sample single cell intensities with a desired CV. The CV was sampled between 0 and 0.3. Synthetic images from these colonies were generated with 3D convolution in the same manner as described in the previous section, but with multiple PSFs: 1) a theoretical PSF being rendered for a 1.49NA objective, a refractive index of 1.518, a working distance of 0.17 mm, and with imaging wavelengths of 0.4 μm, 0.55 μm, and 0.7 μm for separate simulations. 2) the same ground-truth data were convolved with an instrumental PSF captured from a Nikon 100x 1.49NA Plan Apo Ph3 objective lens with the same parameters described, imaged at 0.558 μm wavelength light (bandwidth 40 nm), and 3) the effective PSF fit of the instrumental PSF to simulate long range diffraction effects. This generated synthetic microscopy images and corresponding ground-truth masks of microcolonies under varied imaging conditions. More details on simulation and examples can be seen in supplementary video 1.

#### Colony image intensity quantification and analysis

##### Quantification of single cell intensities from synthetic microcolonies

Calculating the intensity of each cell within the synthetic microcolony images did not require segmentation because the ground truth mask positions are available from the simulations. Thus, for each ground truth mask, the average intensity in the corresponding position in the synthetic microscope image was enumerated and used to calculate the CV, which could be compared to the ground truth CV. To assess whether the ground truth CV can be recovered under ideal circumstances, we performed Richardson-Lucy deconvolution (with 100 iterations) using the original PSF^56,57^. The deconvolved CV can then be compared to the ground truth CV by enumerating the deconvolved image’s average intensity within each ground truth mask position.

The distance of each cell from the centre of the colony was calculated by calculating each cell’s Euclidean distance to the mean position of all cells within the colony (the colony centroid). The distance was normalised by dividing by the maximum Feret’s radius of the colony as calculated by Scikit-image’s regionprops function^58^. The number of neighbours for each cell was calculated by first dilating all cell masks by 4 pixels to ensure that neighbouring cell masks touch. The mask image was then converted into a region adjacency graph, where each node is a cell, and edges represent a neighbour connection between two cells (cells that touch). The graph is then traversed and each node’s degree (corresponding to that cell’s neighbour number) is enumerated (Fig. 5c).

##### Quantification of single cell intensities from experimental microcolonies

Experimental microcolony data were analysed using the same methods as synthetic microcolonies, but were first segmented in phase contrast using the Omnipose model to generate masks (example given in Supplementary Information 24 since only average intensity per cell is required, it is not critical to have very accurate size estimation). Datasets generated this way have no ground truth intensity estimation. Fluorescence images were first background subtracted by subtracting the mean of the lowest 5% of pixel intensities within the image. Mean cell intensity was defined as the sum of the pixel intensities within each mask divided by the cell mask area. Deconvolution was performed using the iPSF and the ePSF, and the intensity of deconvolved cells recorded. Since no ground truth data exists, we did not estimate the CV of real data, but rather focussed on showing the effects of colony size, cell neighbour number, and cell position within a colony on the observed cell intensity. These values were quantified using the same methods as for synthetic colonies.

##### Manual annotation platform and analysis

To assess the effects of projection and diffraction on the performance of manual (human) annotation of cells (for the purposes of training data preparation), five individuals were each asked to annotate the same dataset of 600 simulated cells, where the ground truth was known but not disclosed to the annotators. The cells had an average width of 1 µm, and were partitioned into 4 possible groups: projection “off” cells (cells where the imaging DoF is 1 pixel wide)/projection “on” cells (where the depth of field contains the entire 3D cell), and cells imaged with a fluorescent emitter capable of emitting 0.2 µm wavelength light, and a 0.6 µm wavelength light emitter, distributed with a density of 0.4 emitters per volume element within each cell. 150 of each cell type were scattered with uniform random position and orientation on a 2048×2048 plane with a pixel size of 0.065. Convolution for the two wavelengths was performed once again with the Aguet PSF model. Camera noise was added after convolution using SyMBac’s camera class, using a baseline intensity of 100, a sensitivity of 2.9 analog-digital-units/electron, and a dark noise variance of 8. Annotators ran a Python script which presented them with a Napari window^59^ and a layer upon which to draw on cells. The order in which the images with various effects were displayed was randomised.

##### Deep-learning models for image segmentation

In addition to comparing human annotation accuracy, we sought to test the accuracy of a pretrained model (in this case Omnipose’s bact_fluor_omni model) on simulated images where the perfect ground truth dimension of each cell is known. We generated 200 images containing on average 200 synthetic cells per image (approximately 40,000 total cells) according to the same method described in the previous section, but with areas of synthetic cells varying between 0.15 µm^2^ to 3.5 µm^2^. The PSF model used an imaging wavelength of 600 nm. Ground truth mask and image pairs were saved. Images were then segmented according to the recommended Omnipose documentation with the bact_fluor_omni model. Ground truth cells were matched with their prediction and the IoU for each cell calculated.

We then assessed the benefit of training a model on synthetic data matching one’s experimental setup exactly. The performance gained by training a model on an independent set of synthetic images with perfect ground truth available was checked by generating a test set of 200 images according to the aforementioned method. These were used to train a new model according to the Omnipose documentation for 4000 epochs. The 4000th epoch model was then used to segment the original test data, and ground truth cells were matched with their prediction and IoUs calculated once again.

##### Analysis of cell wall labelled cells on agar pads

After acquisition of cells labelled with FDAAs, images were segmented with Omnipose. Since size estimation of the cell by segmentation is not important, the pre-trained bact_phase_omni model was sufficient to segment the cells. To ensure that all the signal from the fluorescently labelled cell wall was captured, cell masks were binary dilated. After this, all individual cells were cropped, and Scikit-image’s regionprops function^58^ was used to calculate the orientation of the cells and rotate them

##### Simulations of fluorescence images of individual molecules within cells

Fluorescent single molecule images of single cells were generated in the same manner as described before, but with very few emitters per cell (1-30). These low density fluorescent cells were convolved with the tPSF (to capture high frequency information, since long range effects are not needed for this analysis) using the layer-by-layer technique previously described. All analyses were performed on 3 cell types: 1) A typical 1 µm wide, 1 µm deep, 5 µm long cell, 2) An enlarged 2 µm wide, 2 µm deep, 5 µm long cell, 3) A cell trapped by the MACS^10^ platform, 2 µm wide, 0.6 µm deep, 5.5 µm long.

We employed two techniques to count the single molecules in these cells. The first approach, which we term the naive approach, involved sampling fluorescent emitters within the cell and partitioning the emitters into 3 groups: 1) Molecules lost to depth of field; these were defined as molecules more than 0.25 µm away from the centre of the cell. 2) Molecules lost to diffraction; these were defined as molecules residing within 1 Rayleigh diffraction limit of at least one other molecule (with a modification term for the defocus, approximated by the model for the broadening of a Gaussian beam^60^). 3) Resolved molecules; these are the sum of any remaining resolvable single molecules and clusters of molecules within 1 Rayleigh diffraction limit of another (appearing as a single molecule). Rather than applying image processing techniques to count spots, this approach allowed us to identify and partition different sources of miscounting error.

The second approach we applied was Deep-STORM^50^, a deep learning method for super resolution single molecule localisation. A more sophisticated method such as this should perform better than the naive method since it can learn to use defocus, changes in local intensity, and local spatial patterning information to better estimate the number of molecules in a region. We trained Deep-STORM by downloading and modifying the ZeroCostDL4Mic^61^ implementation for local use. While Deep-STORM is typically trained on simulated data, and comes with its own simulator, it does not take into account thick samples such as the depth of entire bacterial cells where defocus is appreciable. Therefore, we generated our own synthetic training data by reducing the number of fluorescent emitters in each cell to between 1-30. Individual models were trained for the regular cell and the MACS cell, both with SNRs (signal to noise ratios) of 8, which is typical for a bacterial single molecule experiment.

All image simulation and image analysis methods made heavy use of scikit-image, NumPy, CuPy, and SciPy.^58,62–64^

### Methods for Experimental Validation

#### Strain preparation

For imaging cells labelled with membrane stains, we used the strain *E. coli* MG1655 7740 ΔmotA with no fluorescent reporter. Cells were grown overnight from a glycerol stock in LB medium at 37°C with 250 RPM shaking. The following day, cells were diluted by 100x into 1 mL of fresh LB. The fresh LB was supplemented simultaneously with both HADA and RADA to a final concentration of 1 mM each. HADA and RADA get incorporated into the bacterial cell wall during growth, allowing imaging of only the cell outline using fluorescence microscopy^65,66^. Cells were allowed to grow in the presence of the FDAAs for 2 hours, after which a 300 µL aliquot was spun down and washed with phosphate buffered saline (PBS) according to the protocol in^67^, taking care to ensure that cells were kept on ice between washes and when not being used.

For imaging microcolonies of fluorescently tagged cells, we used the strain *E. coli* MG1655 7740 ΔmotA with constitutively produced cyan fluorescent protein (SCFP3A, FPbase ID: HFE84) under the control of prpsL. Cells were grown overnight from a glycerol stock in LB medium at 37°C. The following day, cells were diluted by 200x in fresh LB and grown to an OD or 0.1-0.2 to ensure large cell size. Once the desired OD was reached, 1mL of cells were spun down at 4000x g for 5 minutes, and the pellet resuspended in PBS for imaging.

#### Single cell imaging on agar pad

Agar pads were prepared according to the protocol described in Young et al. 2012^27^. Since only snapshot microscopy was to be performed, agar pads were instead prepared with PBS instead of growth medium, and were kept as consistently thick as possible. Agar pads were cut to approximately 22 x 22mm, and placed upon a 22 x 40mm coverslip. Cells on the agar pad were imaged using a Nikon ECLIPSE Ti2 inverted microscope using a 100x (NA=1.49) objective with F-type immersion oil (n=1.518) with a second 1.5x post-magnification lens inserted, for an effective magnification of 150x. The camera used was an ORCA-Fusion C14440-20UP digital CMOS camera from Hamamatsu, with a pixel size of 6.5 µm x 6.5 µm. Cells stained with HADA were imaged with excitation light: 365 nm, 435 nm filter, 100% power, 1 second exposure. RADA was imaged with excitation light: 561 nm, 595 nm filter, 100% power, 0.5 second exposure. Focussing and field of view selection was again done using phase contrast, but special care was taken to account for chromatic aberration by adjusting the Z-plane offset between the focussed phase contrast image and the RADA and HADA images. This was crucial to ensuring that the cell wall was in focus in each image.

#### Imaging microcolonies on agar pads

Since, cells can change their intensities during growth on agar pad, we image preformed colonies to image the effects of diffraction on the cells (example images shown in Supplementary Information 11 and Supplementary Information 15). In order to generate preformed microcolonies, a higher OD of cells (0.1-0.2) was preferred. 3 µL of cell suspension was pipetted directly onto the agar pad and allowed to “dry” for 5 minutes, after which a second 22 x 40mm coverslip was placed upon it. Agar pads were then immediately imaged using the ECLIPSE Ti2 inverted microscope using a 100x (NA=1.49) objective. This enabled us to collect samples of cell clusters (preformed colonies) of various sizes. To avoid photobleaching, well separated fields of view were first selected and focussed in phase contrast. Fluorescent images were captured by excitation with 440 nm light with an LED power of 50% for 900 ms (light source: Lumencore Spectra III Light Engine), and with a filter wavelength of 475 nm. Images were captured as multidimensional 16-bit ND2 files for further analysis.

#### Imaging cells in the mother machine

The mother machine chips were prepared and loaded with cells according to the protocol described in Bakshi et al. 2021^46^. A single mother machine lane was supplied fresh LB by a syringe pump at 15 ul/min. Cells in the mother machine were imaged using the Nikon ECLIPSE Ti2 inverted microscope with a 40x (NA=0.95) objective lens with a 1.5x post-objective magnification lens. The timelapse images were acquired using the Hamamatsu ORCA-Fusion Digital CMOS camera, with a pixel size of 6.5 μm x 6.5 μm. Samples were illuminated with a brightfield light source and a fluorescence light source (Lumencor Spectra III) at 3 minute intervals for 5 hours. Fluorescence images were captured at fast scan mode with a 594 nm excitation LED at 100% power for 100 ms exposure time, and a 632 nm filter.

#### Point spread function acquisition

Our microscope’s (a Nikon ECLIPSE Ti2) point spread function was captured using fluorescent 0.1 µm TetraSpeck Microspheres from Invitrogen. Slides with fluorescent microspheres were prepared according to^68^, with the only change being a bead dilution of 1000x, and the use of Fluoromount-G Mounting Medium from Invitrogen. PSFs were captured using 0.70 NA, 0.95 NA, and 1.49 NA objective lenses, with magnifications of 20x, 40x, and 100x respectively. PSFs were captured with and without the addition of a 1.5x post-magnification lens. Z-stacks were taken of the beads with 0.05 µm spacing. The most in-focus Z stack was determined by taking the radial profile of the PSF and finding the Z-slice with the highest peak intensity and narrowest FWHM. Intensity peaks were then found and beads were selected to maximise the crop area. Bead stacks were then centred around the mean peak intensity and averaged to produce a low noise iPSF.

## Abbreviations

PSF: point spread function
iPSF: instrumental PSF
tPSF: theoretical PSF
ePSF: effective PSF
DoF: Depth of Field
MACS: Microfluidics-Assisted Cell Screening
FDAA: Fluorescent D-Amino Acid
SyMBac: Synthetic Micrographs of Bacteria
FWHM: Full Width at Half Maximum
SNR: Signal-to-Noise Ratio
PSNR: Peak Signal-to-Noise Ratio
CV: Coefficient of Variation
PDMS: Polydimethylsiloxane
HADA: HCC-amino-D-alanine
RADA (also known as TADA): TAMRA 3-amino-d-alanine
IoU: Intersection over Union

## Acknowledgements

We thank Prof. Bartlomiej Waclaw, Prof. Ricardo Henriques, Dr. Diana Fusco, Dr. Temur Yusunov, and Kevin J. Cutler for their feedback on this work and the members of Bakshi Lab for their helpful feedback on this study. All figures included in the present paper are original and contain no third party material.

## Data and code availability

All code written for this paper, and used to generate figures is uploaded to https://github.com/georgeoshardo/projection_diffraction^69^. For backwards compatibility, the version of SyMBac used in this paper has been frozen and included in this repository. Sample datasets, including instrumental point spread functions, microscope images of membrane stained cells and microcolonies, synthetic benchmarking data, and mother machine data have been uploaded to https://zenodo.org/records/10525762^70^.

## Author contributions

G.H. and S.B. conceived of the study. G.H. designed the computational models and deep learning tools. G.H. performed the point spread function, microcolony, and single cell agar pad experiments, and analysed the corresponding data. R.L. performed mother machine experiments and analysis. G.H. and S.B. wrote the manuscript. All co-authors contributed to the final version of the manuscript.

