## Supplementary Information for "Quantitative microbiology with widefield microscopy: navigating optical artefacts for accurate interpretations"

Supplementary Information 1: Image formation in a widefield fluorescence microscope 3
Supplementary Information 2: Additional examples of FDAA-stained cell images 4
Supplementary Information 3: Deconvolution of single cell images fails to correct for the combination of projection and diffraction effects 5
Supplementary Information 4: Effect of depth of DoF and wavelength on cell width estimation 6
Supplementary Information 5: The problem of thresholding cytoplasm images 7
Supplementary Information 6: Diffraction effect on phase-contrast images and choice of emission light filter 8
Supplementary Information 7: An empirical correction factor for membrane width estimation 9
Supplementary Information 8: Benchmarking human annotator performance on simulated data show bias and variability 10
Supplementary Information 9: Performance benchmarking of pretrained Omnipose and SyMBac trained Omnipose. Test data: Synthetic images 11
Supplementary Information 10: Benchmarking performance of pretrained and retrained omnipose on experimental image data 12
Supplementary Information 11: Omnipose retrained on synthetic data outperforms pretrained model on real experiments 15
Supplementary Information 12: Phase retrieval of iPSF 17
Supplementary Information 13: Fitting vectorial model of high NA PSF to the iPSF 18
Supplementary Information 14: Effective PSF (ePSF) to extend the instrumental PSF (iPSF) 19
Supplementary Information 15: Examples of bleedthrough in experimental microcolony images 21
Supplementary Information 16: Additional examples of PSF induced artefacts in gene expression noise estimation 22
Supplementary Information 17: Deconvolution acts as a sharpening filter but does not correct long-range effects in experiments 23
Supplementary Information 18: Long-range effects are not eliminated using small kernel deconvolution 24
Supplementary Information 19: Experiment using a microfluidic device to quantify the extent of bleedthrough in single cell intensity measurements 25
Supplementary Information 20: Multi-snapshot single molecule counting 26
Supplementary Information 21: Deriving a correction factor for depth dependent loss of molecules 27
Supplementary Information 22: Image simulation pipeline 29
Supplementary Information 23: Plane-by-plane convolution is necessary for accurate image simulation 30
Supplementary Information 24: Segmentation of microcolonies for extracting single-cell fluorescence data 32

### Supplementary Information 1: Image formation in a widefield fluorescence microscope

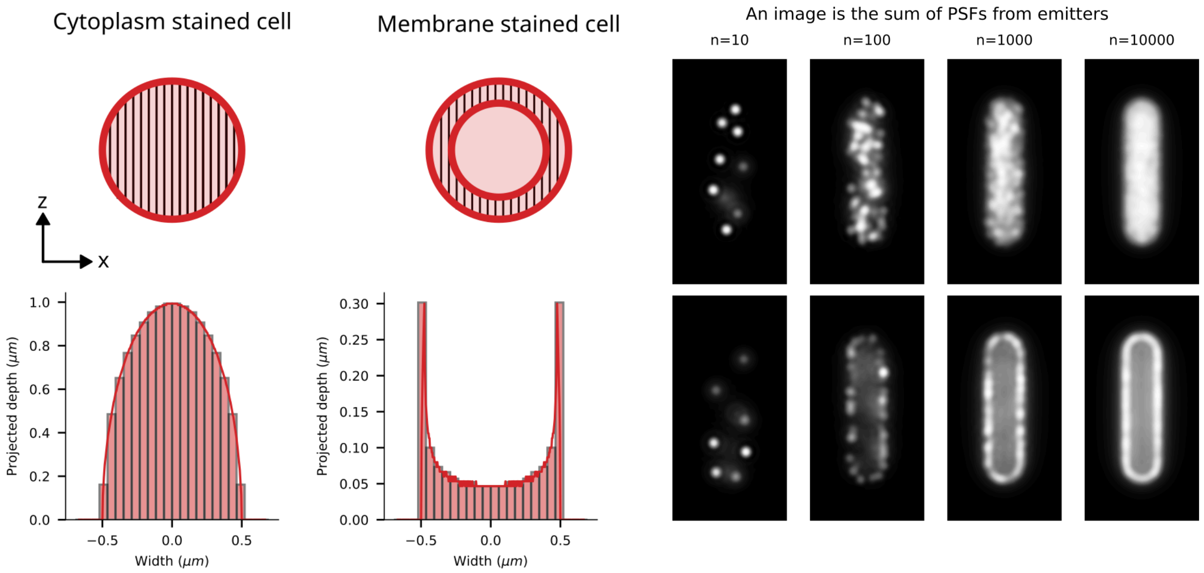


**Figure S1: left)** In 2D widefield microscopy, the projection of volumes along the Z axis causes the intensity profile to appear peaked at the centre for uniformly filled cells and peaked at the edges for membrane-stained cells. **right)** Images taken using a widefield fluorescence microscope are made up of point sources of light. Each point source of light is imaged as a point spread function (PSF). The exact shape of the PSF is determined by the optics used to image the point source, and will be unique to each imaging system. The shape of the PSF is also variable dependent on the position of the point emitter in the Z plane. Thus, an image taken using a widefield fluorescence microscope is the sum of many point spread functions. Two examples are shown: top - cytoplasm uniformly filled with emitters, bottom - emitters placed on membrane.

### Supplementary Information 2: Additional examples of FDAA-stained cell images

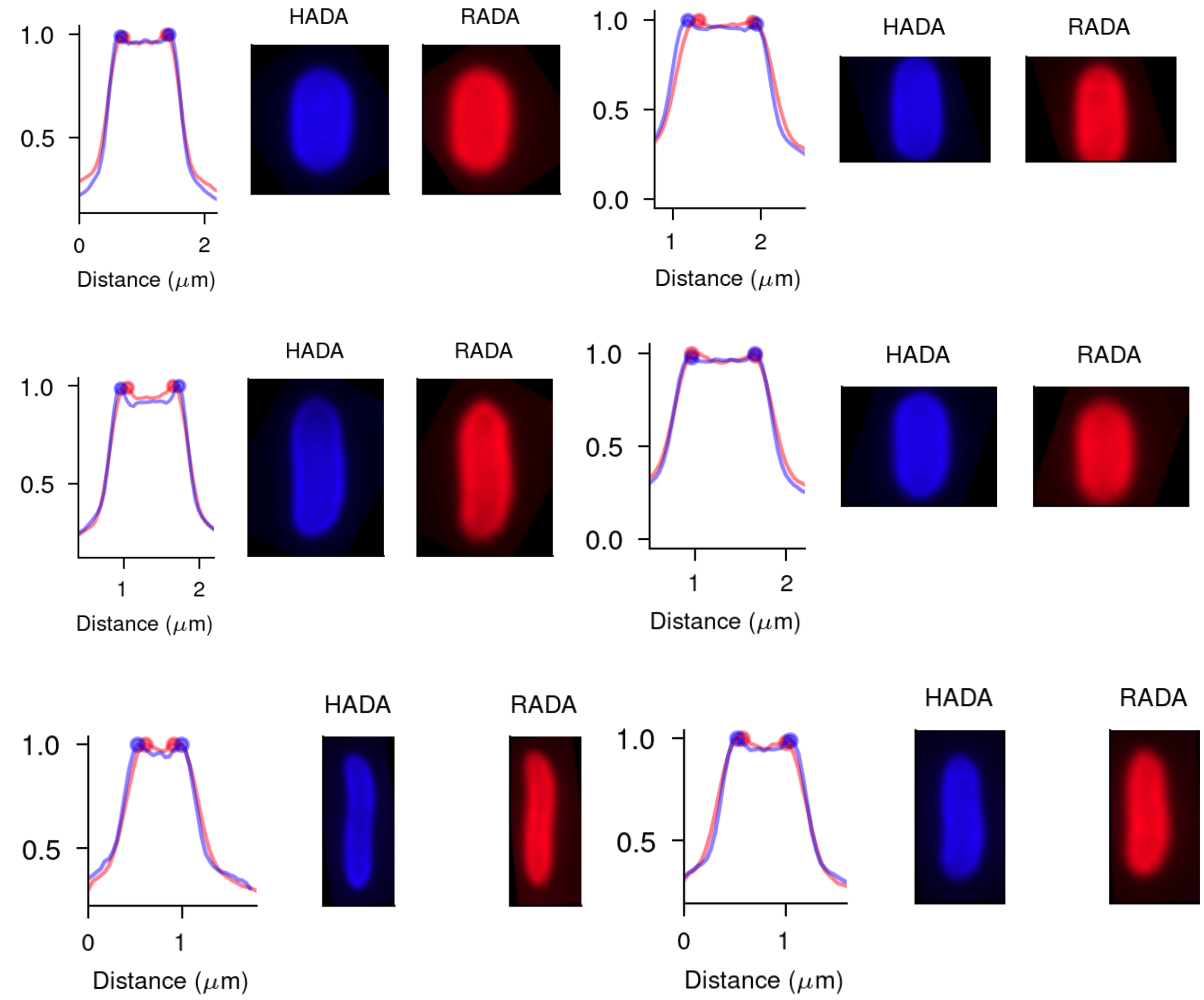


**Figure S2:** Example snapshots of cells imaged with both blue FDAA (HADA) and orange FDAA (RADA) and the corresponding radial intensity profiles in respective colour channels. Due to diffraction effects, red traces are always wider at the tail and narrower at the peaks (circles).

#### Supplementary Information 3: Deconvolution of single cell images fails to correct for the combination of projection and diffraction effects

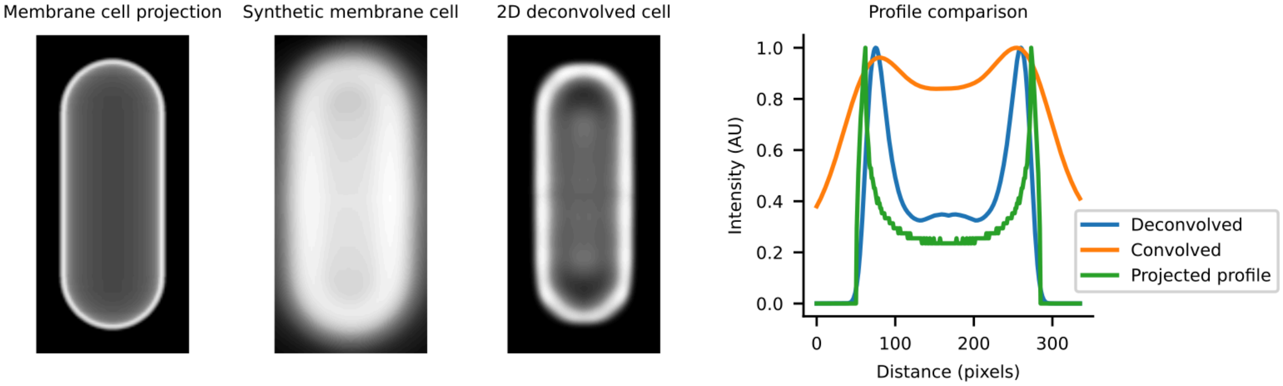


**Figure S3:** The result of 2D Richardson-Lucy deconvolution (with 200 iterations) of a membrane stained cell. Deconvolution cannot fully reconstruct the image (as can be seen in the profile comparison) due to the effects of projection. It simply results in a sharper profile, with its peak located in the same erroneous location. Increased iterations of deconvolution does not increase the profile reconstruction accuracy, and instead result in deconvolution ringing around the centre of the cell.

### Supplementary Information 4: Effect of depth of DoF and wavelength on cell width estimation

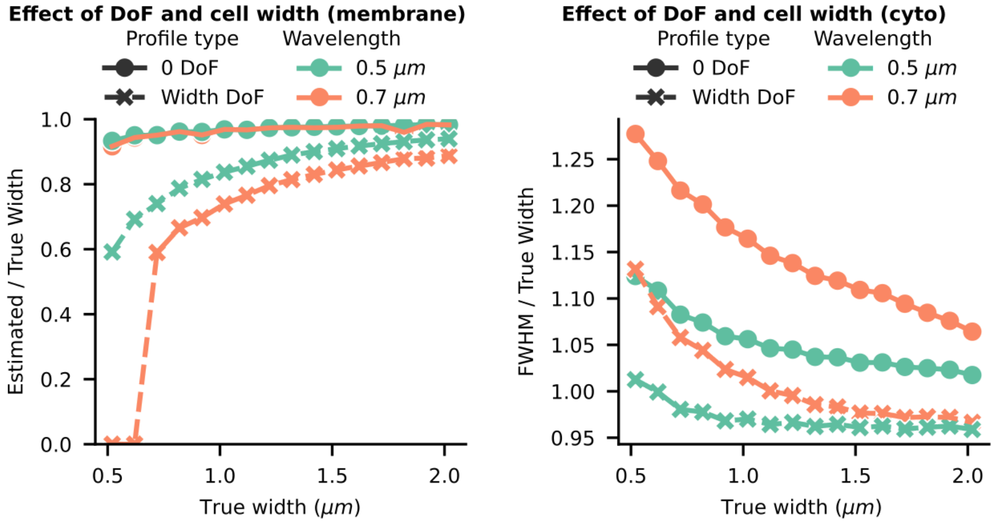


**Figure S4:** Trendlines from synthetic data show that the ratio of estimated width and true width is dependent on the cell width for both membrane labelled cells (left), and for cytoplasm stained cells (right). The error grows rapidly for narrow cells and for cells with widths below 600 nm, the membrane stain method fails to work. The depth of field (DoF) = 0 lines show that without projection, the effects of diffraction and cell-width dependence of the results are mitigated for membrane stained cells. However, DoF=0 increases the error for cytoplasmically labelled cells. The estimated widths of cytoplasmically stained cells are calculated by taking the full width half maximum (FWHM) of the radial profile of the intensity, whereas for membrane stained cells the interpeak distance is taken.

#### Supplementary Information 5: The problem of thresholding cytoplasm images

Thresholding cytoplasmically fluorescent cells poses a problem. Unlike with membrane stained cells, there is no parameter such as interpeak distance which can be quantified without bias. An image like this is blurred, and the blurring profile is imaging system dependent. Drawing a threshold around a blurred image is subject to bias, from the imaging wavelength, imaging system, and the thresholding method.

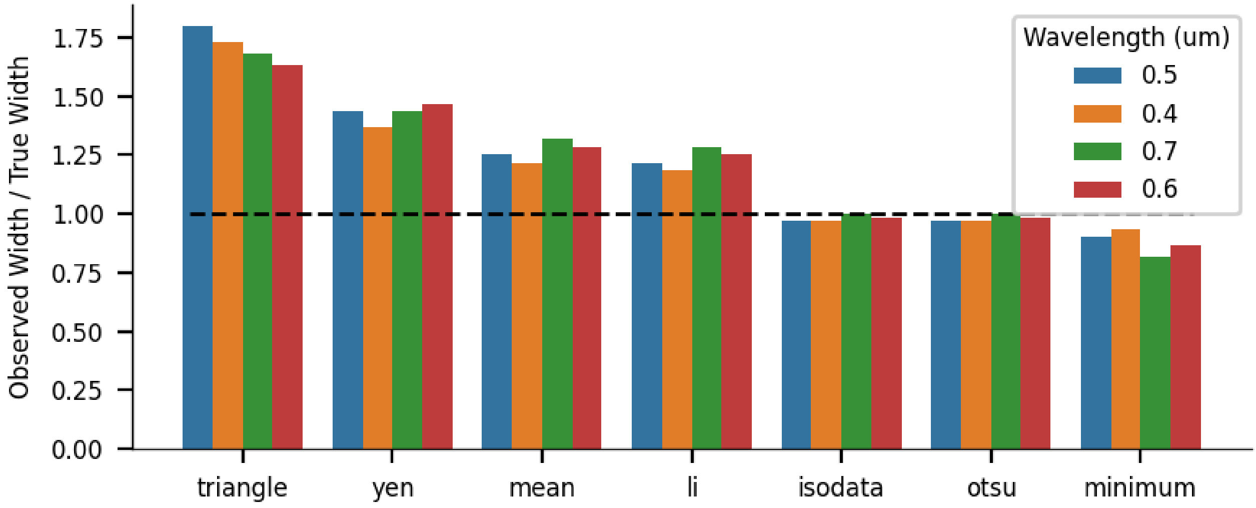


**Figure S5**: The quality of output masks from various thresholding algorithms are compared. A 1 μm wide, 3 μm long synthetic cytoplasmically fluorescent cell is simulated under various emission wavelengths (1.49NA, 1.51 refractive index), and thresholded with common global thresholding techniques. Most thresholding algorithms show inaccurate and wavelength dependent performance, except isodata and otsu.

### Supplementary Information 6: Diffraction effect on phase-contrast images and choice of emission light filter

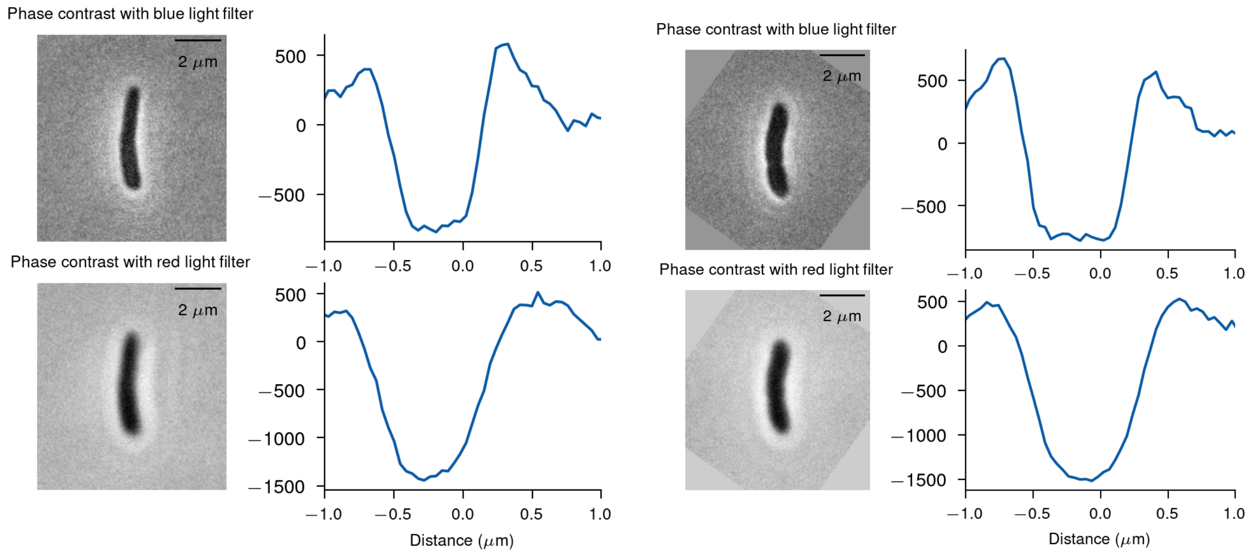


**Figure S6:** Example phase-contrast images of the same cells collected with different emission filters (blue - 435 nm and red 595 nm). Images formed using the lower wavelength light are sharper and produce narrower cross sections (as seen from the corresponding radial intensity profiles), while images from longer wavelength are washed out due to diffraction effects. Note the flatness of the minima of the blue light profiles in comparison to the red light profiles. This is likely due to the narrower z-profile of the transmitted phase contrast PSF under blue light conditions (effectively resulting in less contribution from projected planes and more from the central plane).

### Supplementary Information 7: An empirical correction factor for membrane width estimation

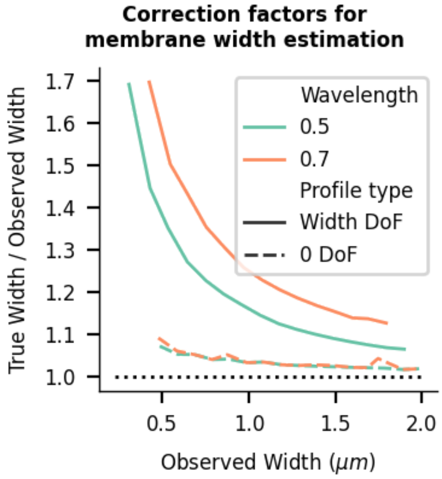


**Figure S7**: Virtual microscopy can be used to estimate correction factors for observed cell widths in diffraction limited scenarios, given that the user's imaging system is well modelled. The correction factor is close to 1, when the depth of field is very shallow.

#### Supplementary Information 8: Benchmarking human annotator performance on simulated data show bias and variability

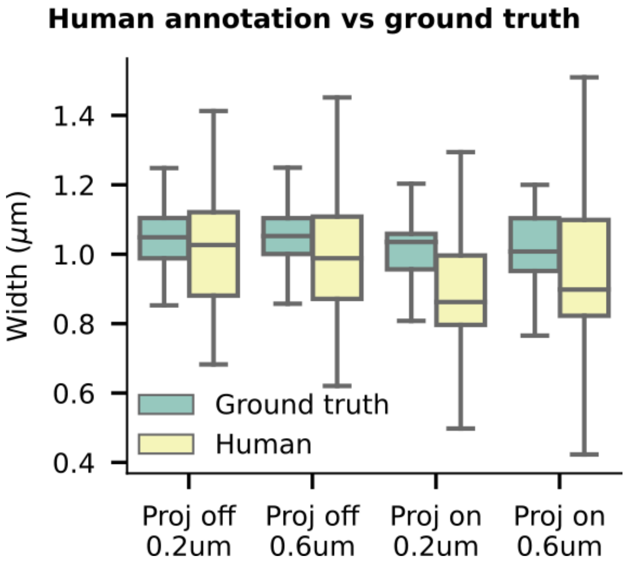


**Figure S8**: Synthetic data of cytoplasmically stained cells is simulated to test human annotation (1.49NA, 1.51 refractive index, 0.065 μm/pixel). To test whether annotation improved when there was almost no diffraction, we also convolved some of the data with a theoretical 0.2 μm PSF, as well as a 0.6 μm PSF. Additionally cells were simulated with no projection. Human annotation performance on such datasets is inconsistent, both underestimating the mean (most likely due to projection effects), and having very high variability.

### Supplementary Information 9: Performance benchmarking of pretrained Omnipose and SyMBac trained Omnipose. Test data: Synthetic images

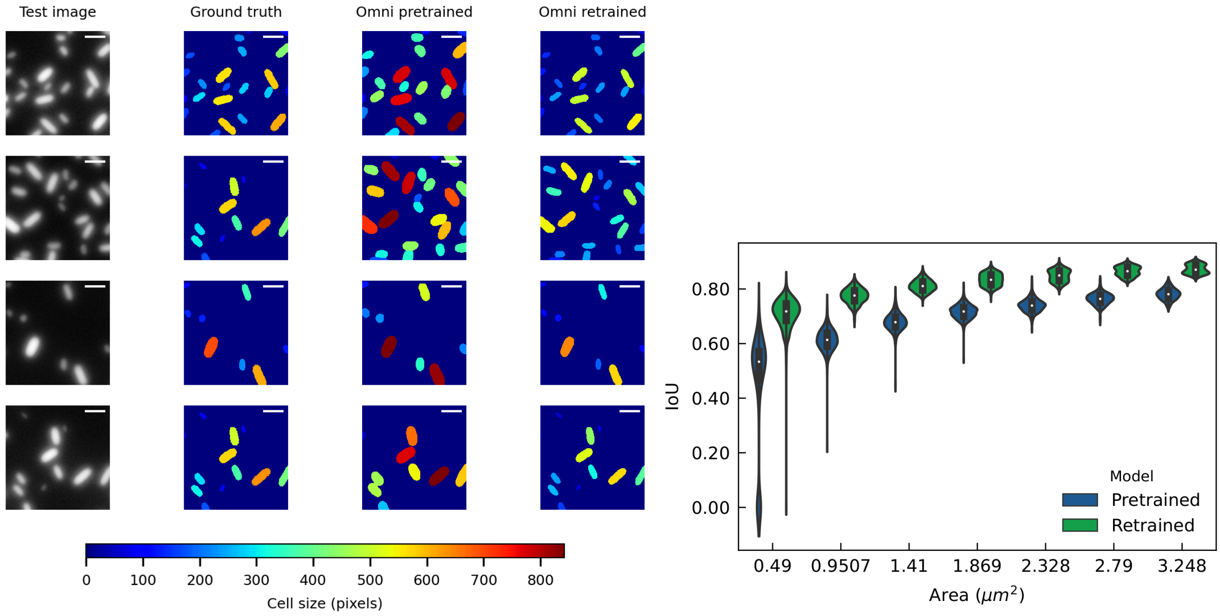


**Figure S9: left)** Comparison of segmentation performance of the pretrained Omnipose model (bact_fluor_omni) and the Omnipose model retrained with synthetic data that closely matches the target data. Example images of the test sample are shown on the left column and the corresponding ground truth masks are shown on the column next to it. The segmentation output from the pretrained Omnipose model (trained with human annotated data) is shown on the next column (third from the left). Corresponding segmentation outputs from the omnipose model retrained with synthetic data are shown on the rightmost column. To compare the segmentation results and ground truth, each mask is coloured according to its size (scale bar shown at the bottom). Close correspondence to the ground truth and robust performance across cell size and intensity variation is noticeable. **right)** The retrained models improve the performance in terms of segmentation accuracy and precision, as shown by increased IoUs and reduced variabilities for various sizes of the digital cells. The performance in each cell area class can be further boosted by training bespoke models for each cell size range.

#### Supplementary Information 10: Benchmarking performance of pretrained and retrained omnipose on experimental image data

Benchmarking deep learning models on microscope data presents inherent challenges due to the absence of accurate ground truth. The point spread function inherently blurs the image, making it impractical for human annotators to draw accurate masks around objects. Consequently, while it may be relatively straightforward to assess the performance of deep learning models on synthetic test data, given the availability of perfect ground truth, evaluating their performance on real data poses greater difficulty, as perfect ground truth is lacking.

To tackle this challenge, we devised an experiment aimed at benchmarking the performance of segmentation models on real data, extracting the ground truth properties of cells from experimental images. Our method involves culturing dense populations of cells on agarose pads, leading to the formation of instantaneous cell clusters, which we term "preformed colonies." Importantly, the properties of individual cells within these colonies mirror the distribution observed in the original culture, regardless of whether they are clustered or isolated on the pad. Moreover, since bacterial cell width is tightly regulated, we can reasonably assume that the width distribution remains consistent across cells within clusters of varying sizes. By accurately measuring the mean width of cells from aligned patches within these clusters, as illustrated in Fig. S10 below, we can retrieve the ground truth cell width, and effectively benchmark the performance of deep learning segmentation algorithms trained on diverse training data.

Images of preformed bacterial colonies were captured following the protocol outlined in the methods section. We identified clusters containing five or more cells arranged side by side. We infer that this cellular arrangement results from cells pushing on and aligning with one another during colony formation, facilitated by the placement of the coverslip onto the agar pad and spreading of the droplet. Thus, we assume physical contact between these cells. Given the tightly controlled width of E. coli, we can estimate the true mean and standard deviation of cell width by measuring the total width of several patches and dividing by the number of cells within each patch. Subsequently, we manually retrieved stacks of cells from both SyMBac-trained and pretrained Omnipose models. The widths of these stacks were measured, and the width of a single cell was estimated by calculating the mean cell width across each patch and then aggregating the grand mean across all patches. We observed nearly identical mean and standard deviations (see Table S1).

Subsequently, we conducted a comparison between the performance of the Omnipose (bact_fluor_omni) model trained using human-annotated data (pretrained) and the same model retrained with synthetic training data generated by SyMBac (retrained). Example fields of view of the test data, along with their associated segmentation outputs, are presented in Fig. S11. These examples illustrate that the output of the pretrained omnipose model exhibits variability between cells, depending on whether they are isolated or part of clusters, as well as their position within the cluster. In contrast, the retrained Omnipose model demonstrates robust performance, aligning closely with the predictions from the benchmarking analysis conducted on synthetic test data. The mean cell widths obtained from the retrained omnipose model (0.94 ± 0.062 μm) closely correspond to the estimates derived from the patch analysis (0.94 ± 0.064 μm), whereas the output of the original omnipose model tends to overestimate cell width with notable variability (see Table S2).

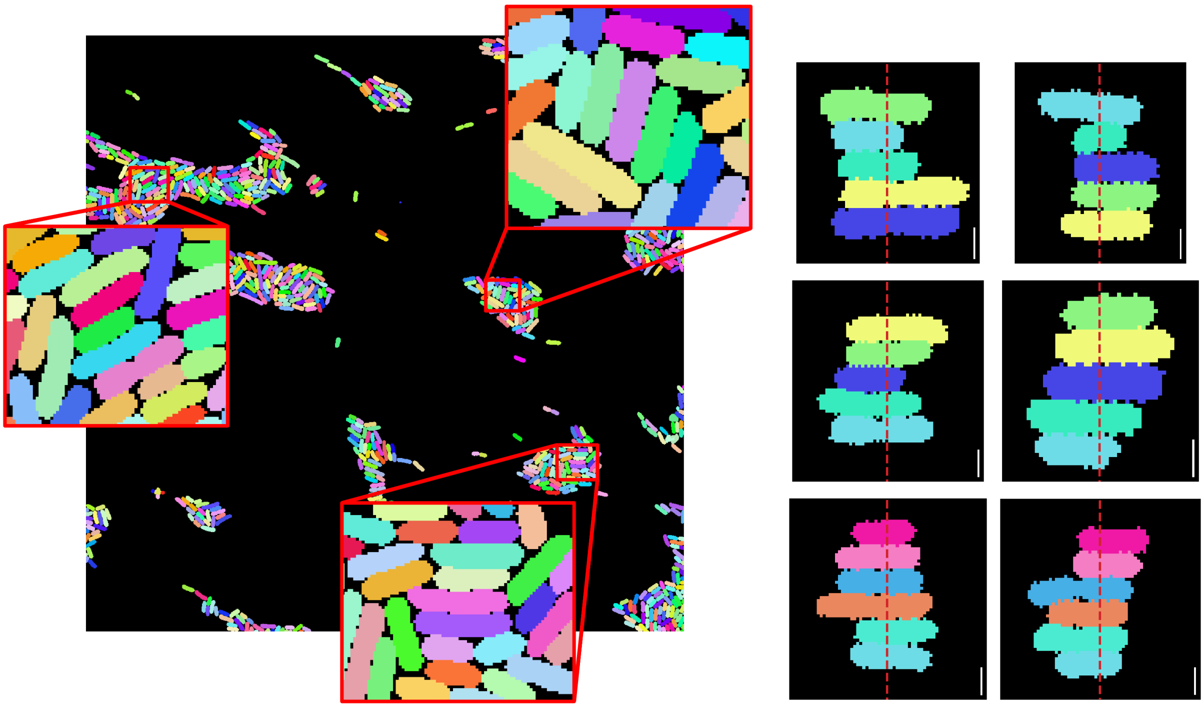

**Figure S10**: **left)** An example field of view from the segmented data, with patches of aligned cells zoomed. **right)** 12 patches were analysed (6 shown), for a total of 66 cells.Their overall width is estimated along the red line. We took the width to be the average of a 4 pixel wide line, to minimise small pixelation artefacts at the edges of cells arising from rotation without pixel interpolation. (Scale bar = 1 μm)

| | Patch mean width per cell | Standard deviation |
|---|---|---|
| SyMBac Omnipose | 0.95 μm | 0.064 μm |
| bact_fluor_omni Omnipose | 0.93 μm | 0.068 μm |
| **Average model statistics** | **0.94 μm** | **0.066 μm** |

**Table S1:** True mean and standard deviation of the width of cells from models. Average between models is taken to be the ground truth. Determining the true mean cell width from patches is not dependent on the model selection, since we chose patches of cells which are tightly packed and space filling, thus masks for cells within the patch necessarily must

#### Supplementary Information 11: Omnipose retrained on synthetic data outperforms pretrained model on real experiments

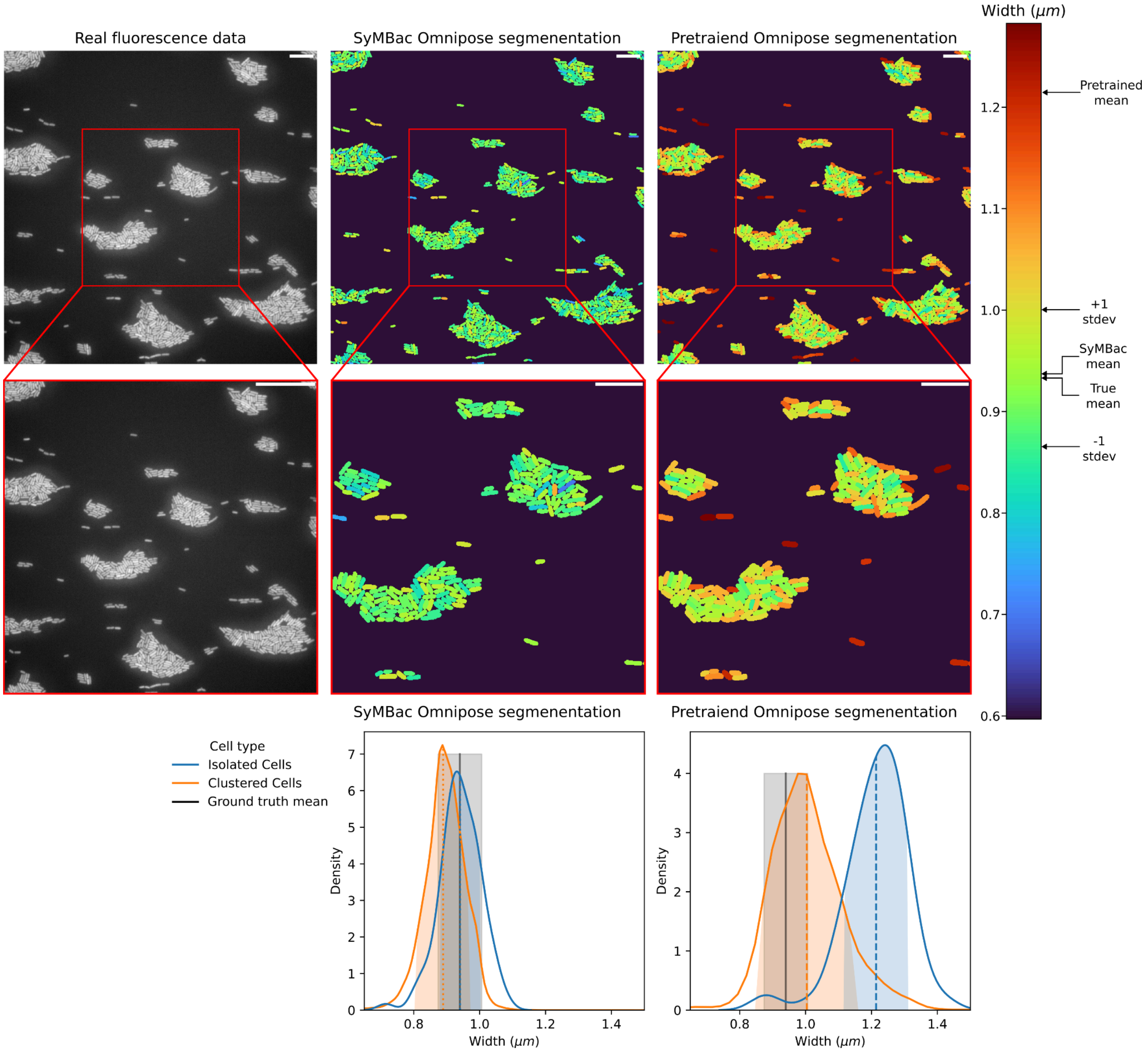


**Figure S11**. Segmentation masks for an example field of view (shown left), from both SyMBac trained (middle) and pretrained (right) Omnipose models. The mask colour is given according to the segmented width of each cell, calculated by taking the number of pixels intersecting a line drawn perpendicular to the cell's long axis. Full population distributions are shown below (Scale bar = 5 µm). The SyMBac retrained model gave robust and accurate performance. The width of the mask was independent of the position of a cell, unlike the pretrained model, which overestimated sizes of isolated cells compared to the cells in clusters, and had larger variability in the output.

| Cell type | Model | Width (μm) | Standard deviation |
| --- | --- | --- | --- |
| Isolated cells<br>N = 95 | SyMBac Omnipose | 0.94 μm | 0.062 μm |
| | bact_fluor_omni | 1.2 μm | 0.10 μm |
| Clustered cells<br>N = 12282 | SyMBac Omnipose | 0.90 μm | 0.082 μm |
| | bact_fluor_omni | 1.0 μm | 0.15 μm |

**Table S2:** Retraining on bespoke synthetic data tailored to the experiment brings the width estimates and standard deviations to well within bounds of the estimated true mean. The retrained model shows higher precision (lower standard deviation) compared to the pretrained model.

### Supplementary Information 12: Phase retrieval of iPSF

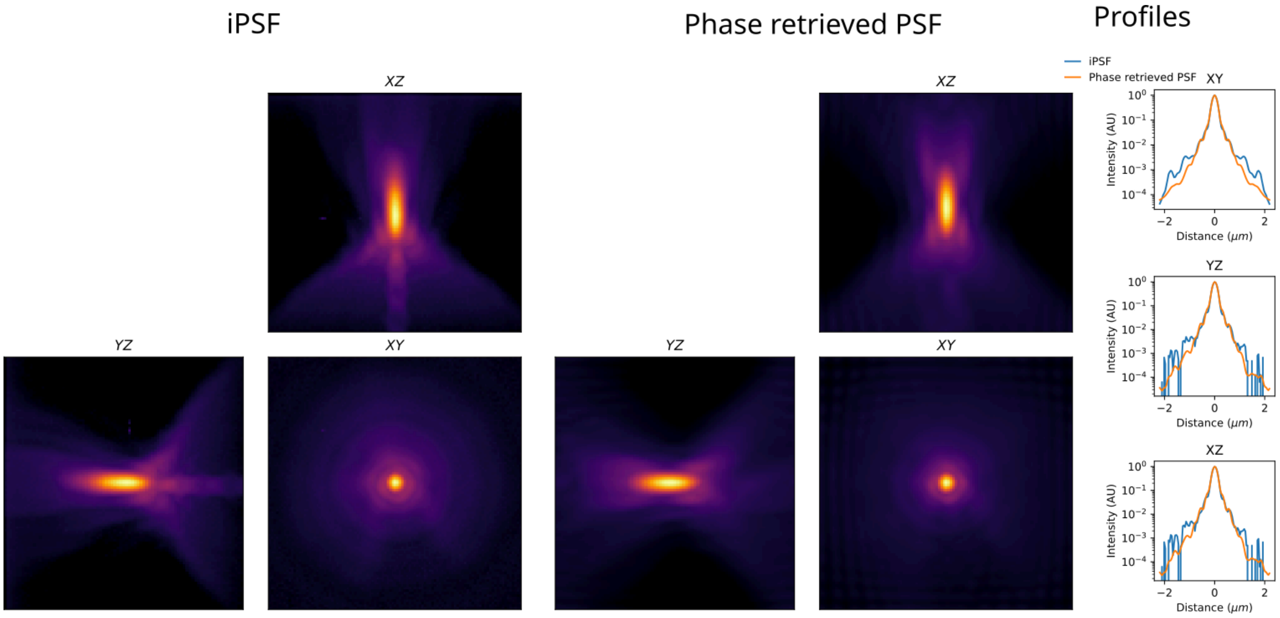


**Figure S12:** Using the Hanser phase retrieval method[1] implementation created in https://github.com/david-hoffman/pyOTF, and manually adding the empirical Gaussian rescaling function to the derived OTF, we were able to fit a phase retrieved PSF to our iPSF, incorporating aberrations within our microscope. However, we found the method to be extremely sensitive to background subtraction (responsible for the noise at the edges of the iPSF in the yz and xz profiles on the right). Additionally, while the extrapolation beyond a +/- 1 μm (shown in the right profiles) was better than solely using the tPSF (presumably due to the addition of fitted aberrations), it was not as performant as using the ePSF. It is possible that with an even higher SNR iPSF, and by considering field-dependent effects, phase retrieved PSFs would be the ideal solution.

### Supplementary Information 13: Fitting vectorial model of high NA PSF to the iPSF

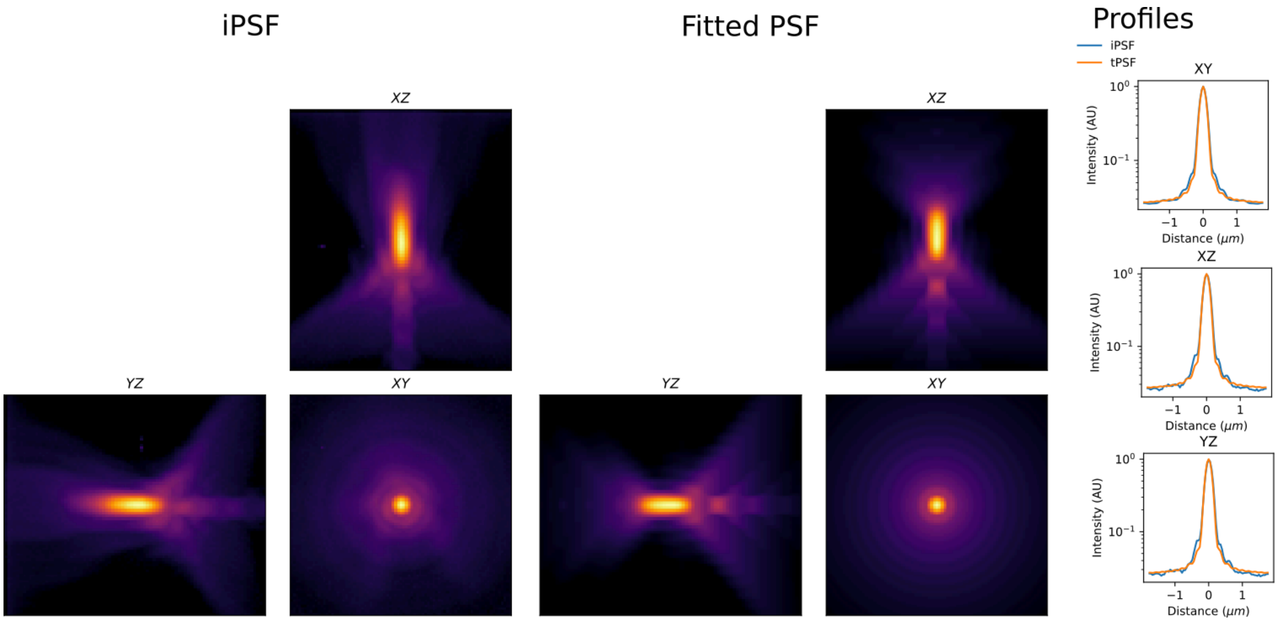


**Figure S13:** The vectorial PSF model from Aguet[2], when fitted to the iPSF (wavelength = 515 nm, NA = 1.45) yields a good fit, and while this can be used for image simulation of small structures, extrapolation of the function's domain beyond the fit yields poor results (as shown in Supplementary Information 14, given the same fitting methodology).

### Supplementary Information 14: Effective PSF (ePSF) to extend the instrumental PSF (iPSF)

For accurate image simulation, convolution must be done with both a kernel which is a good representation of the imaging system's point spread function (PSF), and large enough to capture long range diffraction effects. Since the instrumental PSF (iPSF) can only be captured experimentally for short to medium distances (owing to loss of signal to the noise floor and limited field of view), we sought to "extend" the range of the iPSF by fitting an effective PSF (ePSF) to it. This was done by choosing functions and fitting them to the images of each Z-plane of the iPSF. Once a good fit is found, the x-y domain of the function can then be extrapolated to estimate long range effects. The main assumption of this method is that the local fit of the function to the iPSF will yield an accurate extrapolation beyond the measured x-y domain. For our imaging system, we found that a Lorentzian function modified with a difference of Gaussians yielded a good fit. The model is given by:

$$
\begin{aligned}
\mathcal{L}(x,y) &= \frac{A}{1+\frac{x-x_0}{\gamma_x}^2+\frac{y-y_o}{\gamma_y}^2}+C \\
\mathcal{G}_1(x,y) &= A_{g_1}\exp\left(-\frac{(x-x_0)^2+(y-y0)^2}{2\sigma_1^2}\right)+C \\
\mathcal{G}_2(x,y) &= A_{g_2}\exp\left(-\frac{(x-x_0)^2+(y-y0)^2}{2\sigma_2^2}\right) \\
\mathrm{DoG}(x,y) &= \mathcal{G}_1(x,y)+\mathcal{G}_2(x,y)+\max(0,-\min_{x,y}(\mathcal{G}_1(x,y)+\mathcal{G}_2(x,y)) \\
\mathrm{PSF}(x,y) &= \mathcal{L}(x,y)\cdot\mathrm{DoG}(x,y)
\end{aligned}
$$

Where:

- $A$ is the amplitude of the Lorentzian term.
- $x_0$ and $y_0$ are the centre coordinates of the function.
- $\gamma_x$ and $\gamma_y$ are the Lorentzian scale factors.
- $A_{g_1}$ and $A_{g_2}$ are the amplitudes of the Gaussians.
- $\sigma_1$ and $\sigma_2$ are the Gaussian scale factors.
- $C$ is a small constant offset.

Fitting of the model to the iPSF was set up as a global optimisation problem and performed using differential evolution with the loss function being defined as:

$$
\mathrm{Loss}=\frac{1}{MN}\sum_{i=1}^{M}\sum_{j=1}^{N}\left(\ln(y_{ij}+\epsilon)-\ln(\hat{y}_{ij}+\epsilon)\right)^2
$$

Where:

- $M$ is the height and $N$ is the width of the PSF and the fit.
- $\epsilon$ is a machine epsilon.

This loss function will be large even for differences in small values, which is useful for fitting PSFs, where the magnitude of the values spans 3 or more orders of magnitude. Therefore, the fit of the tails is maintained using this loss function.

.

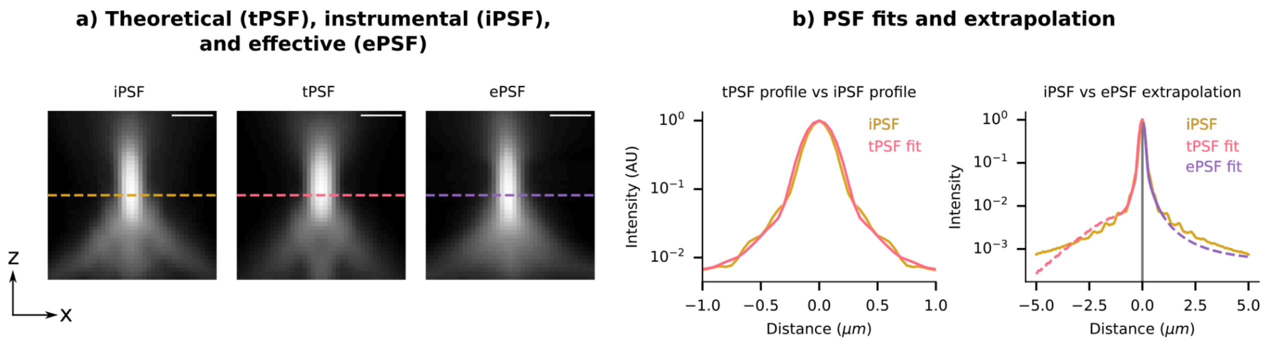


**Figure S14: a)** The instrumental (i)PSF, theoretical (t)PSF, and effective (e)PSFs are all shown in the xz plane. The tPSF and ePSF are both fitted to the iPSF according to the aforementioned method. The dotted line shows the location of the drawn intensity profile in panel b. **b)** The tPSF can be fitted to the iPSF, however for simulation of very long range optical effects, such as diffraction over a long distance within a microcolony, the iPSF would have to be measured with very high SNR up to ranges of more than tens of microns. This would require an extremely large number of very highly separated beads to be imaged and averaged. Additionally, the magnitude of the PSF's intensity may vary over more than 4-5 orders of magnitude as the domain expands, making capturing the entire PSF's range impossible within the values of the 16-bit unsigned integer type, which many microscope cameras use. Therefore we attempted to extrapolate the domain of the tPSF to simulate long range effects. This showed that the theoretical PSF (tPSF) was not a good fit to the iPSF compared to the optimised ePSF function. With ePSF domain extrapolation, one can generate a PSF of any size, and use 64-bits of floating point precision for convolution. However, we note here that the ePSF still under-estimates the long range effects that we measured our iPSF to have at ~5 μm from the centre.

#### Supplementary Information 15: Examples of bleedthrough in experimental microcolony images

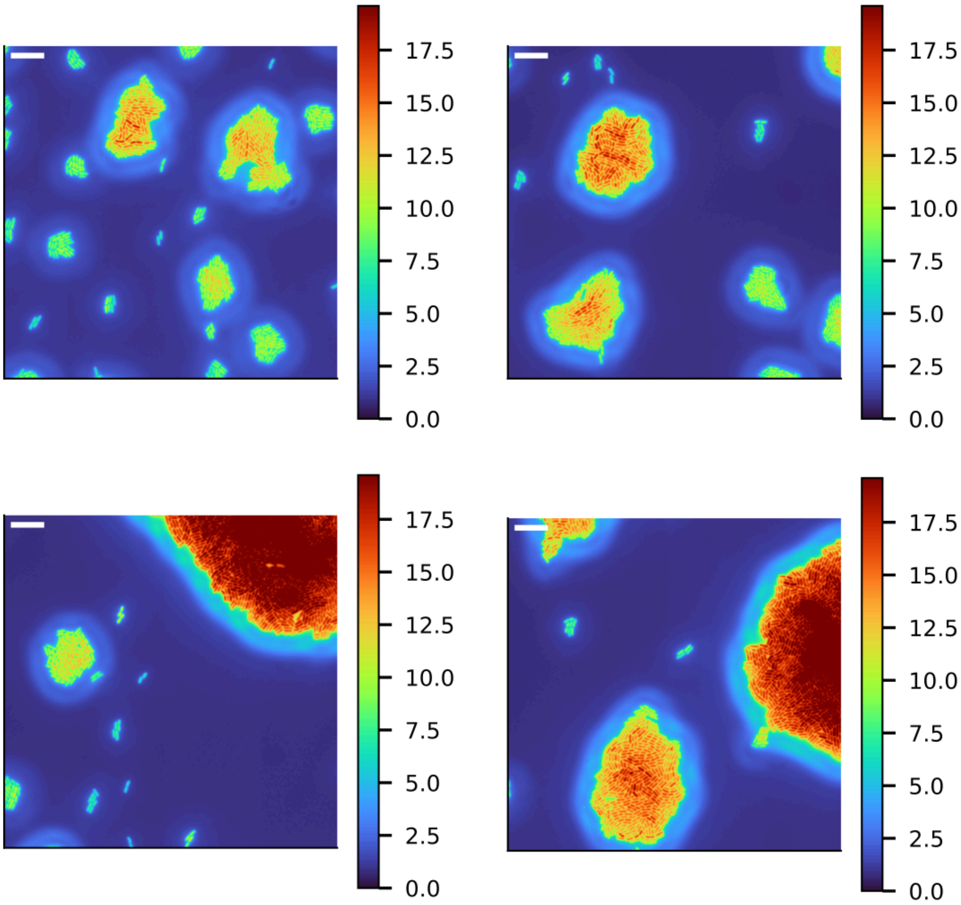


**Figure S15:** Additional examples of experimental microcolonies showing the bleedthrough effect. Isolated cells and cells in smaller colonies appear to have lower intensities than larger microcolonies. Position dependent effect on cells is seen within each colony, as cells near the colony centre appear brighter than cells near the perimeter. (Scale Bar = 10 µm, images captured with 100x Plan Apo oil objective, cells expressing SCFP3A from pRpSL promoter, emission filter 480/40 nm).

#### Supplementary Information 16: Additional examples of PSF induced artefacts in gene expression noise estimation

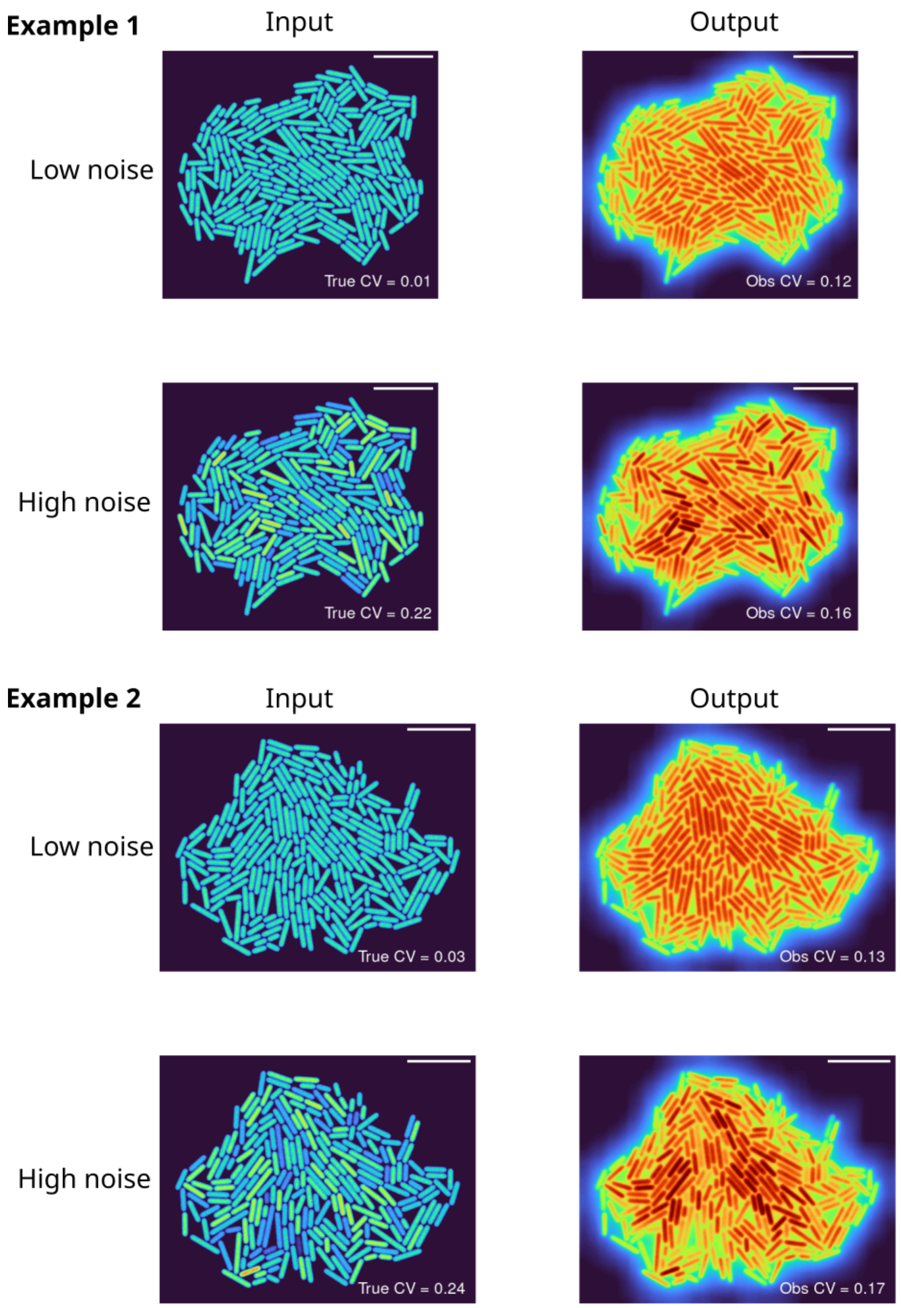


**Figure S16:** Two examples of synthetic images of simulated microcolonies composed of cells with different intensity distributions (low and high coefficient of variation gene expression). Both examples show how convolution with the PSF causes low noise microcolonies to be observed as higher noise than they are, and high noise microcolonies to appear as low-noise.

### Supplementary Information 17: Deconvolution acts as a sharpening filter but does not correct long-range effects in experiments

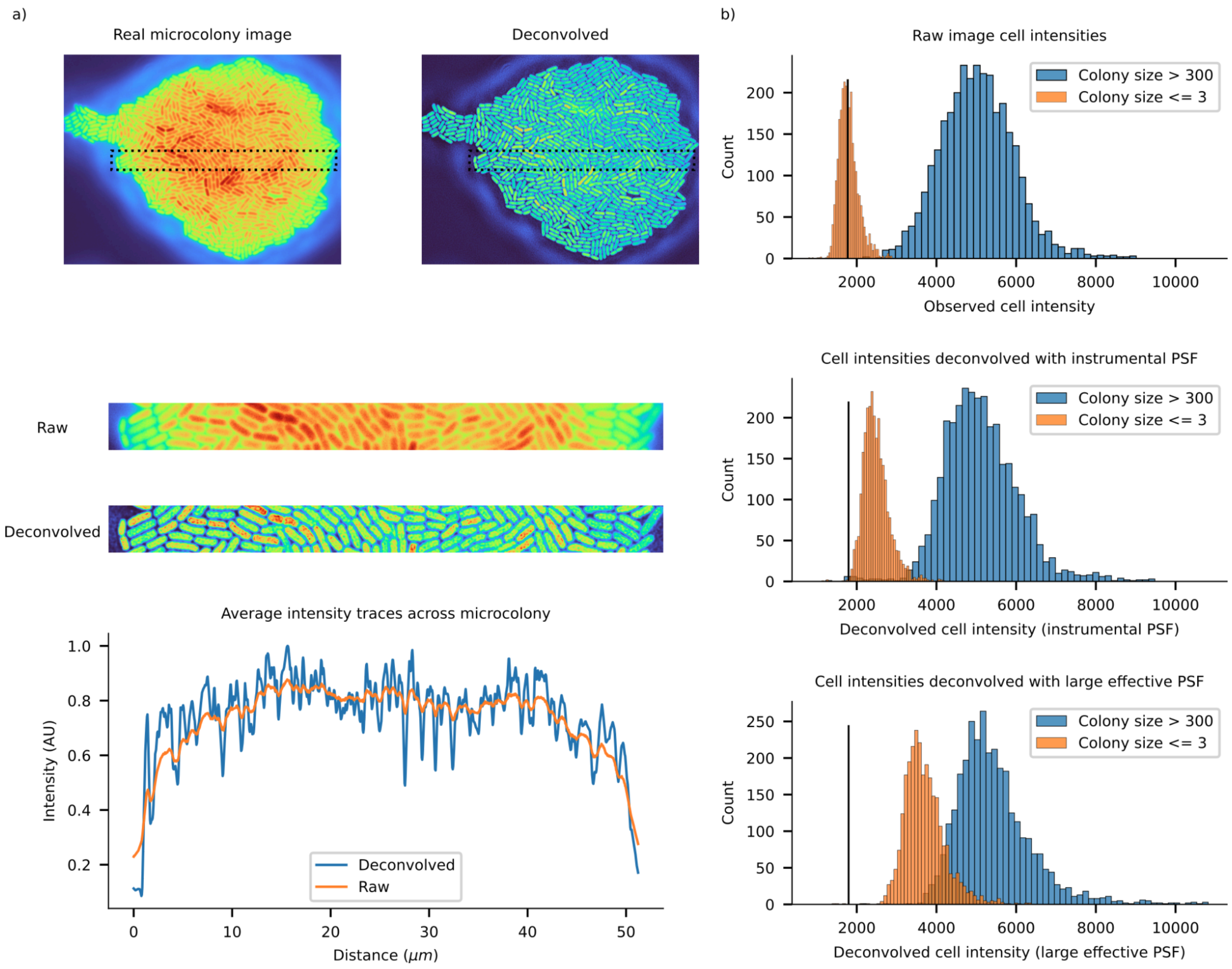


**Figure S17:** Deconvolution of real images using the instrumental PSF acts as a sharpening filter. a) shows a real microcolony image and its deconvolved counterpart (in Turbo colour-map). Initially it appears that the deconvolution has corrected the bright centre of the colony, restoring the correct intensities of all cells. Instead what can be seen from the crops is that the deconvolution has simply sharpened the boundary between individual cells, and an intensity profile across the colony reveals that the overall trend is the same. b) shows that the intensity of cells within large microcolonies remains largely unchanged after deconvolution, while single cells get brighter post-deconvolution. This is because in large colonies, a cell loses intensity to its neighbours, but those neighbours in turn lose intensity to that cell, and the effect is cancelled out. Cells in small colonies lose intensity to the surrounding background, and there are no neighbours to repay the lost intensity. As can be seen in the third panel of b), the correction in single cell intensities is better with large effective PSFs, but is still not capable of completely restoring the image.

### Supplementary Information 18: Long-range effects are not eliminated using small kernel deconvolution

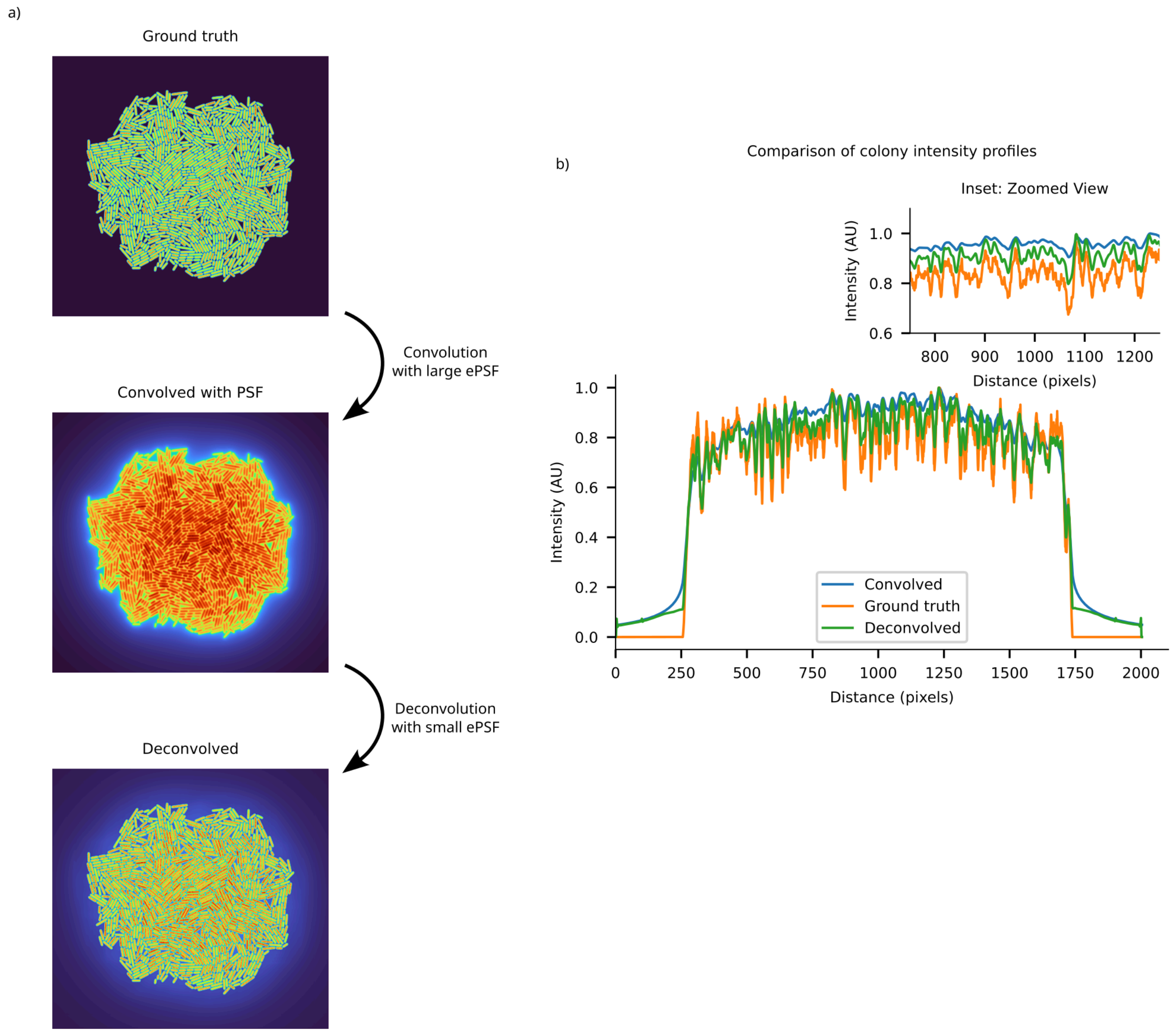


**Figure S18:** a) A synthetic microcolony is convolved with a large effective PSF to simulate image formation, and then deconvolved with a smaller version of the effective PSF to mimic the fundamental limitations of deconvolution using experimentally measured instrumental PSFs. The smaller ePSF is unable to correct for long range effects, and some of the image corruption remains in the form of a halo around the colony and brighter cells in the colony centre. b) Line profiles drawn through the middle centre of the three colony images in panel a. The result of improper deconvolution manifests as only a partially restored line profile.

### Supplementary Information 19: Experiment using a microfluidic device to quantify the extent of bleedthrough in single cell intensity measurements

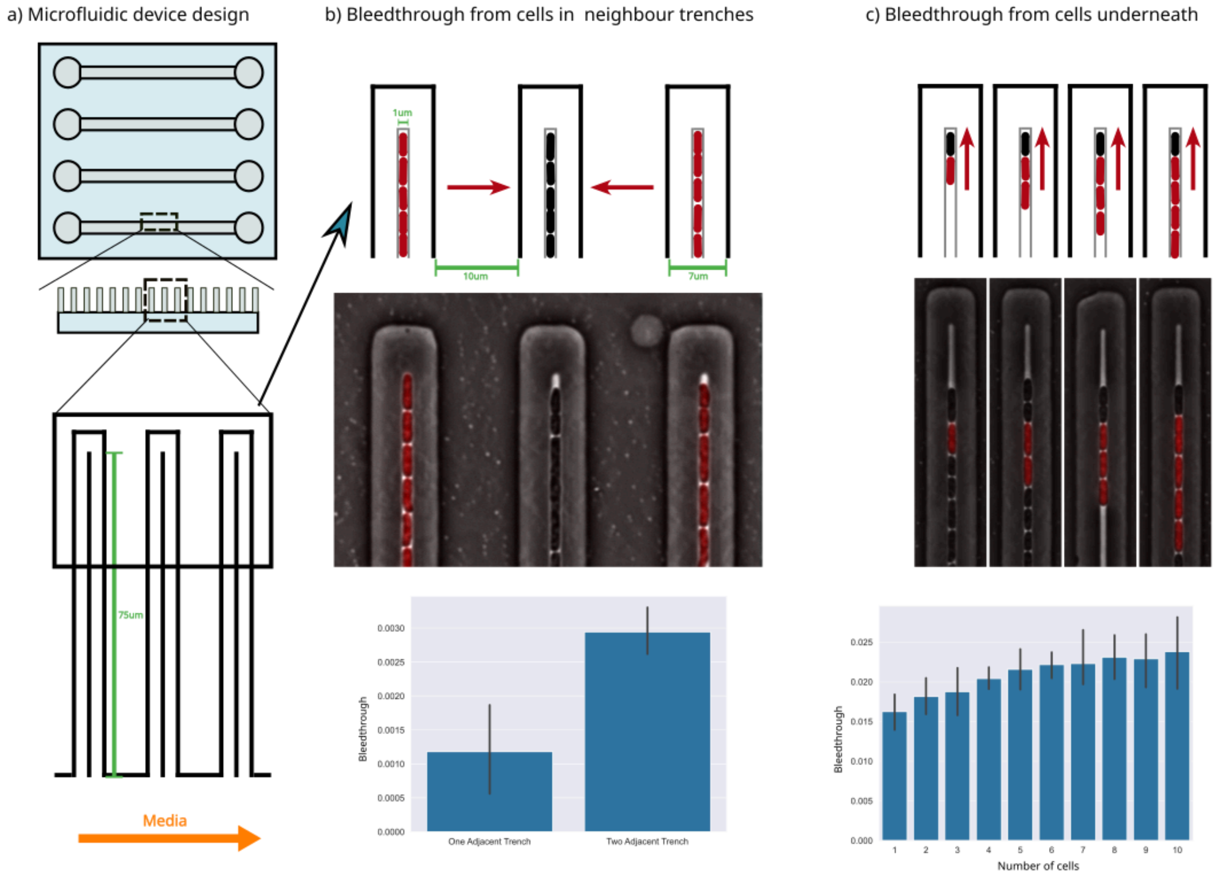


**Figure S19:**. **a)** The schematic depicts the design of the specific microfluidic device known as the 'mother machine.' This device consists of horizontal lanes equipped with inlets and outlets for the flow of growth media. The zoomed version of the inset below shows how narrow trenches (1.5 um wide) are placed orthogonally next to each lane. These trenches comprise two layers: one with a depth of 1 μm to house cells, and another wider and shallower layer (400 nm deep) to deliver media to the cells, as illustrated in the inset at the bottom of this page. **b)** We combined fluorescently labelled (Red) and unlabeled cells (Black) within the device to assess bleedthrough effects from cells in adjacent trenches. The top schematic illustrates the configuration used to quantify bleedthrough from cells in two neighbouring trenches, while a representative snapshot from the experiment is presented below. The data plot depicts the extent of bleedthrough, measured as the estimated intensity of unlabeled cells above background in relation to the number of cells in neighbouring trenches ($bleedthrough = \frac{I_{unlabelled} - I_{background}}{I_{labelled} - I_{background}}$). For cells in one adjacent trench, their contribution to the intensity of unlabeled cells is slightly above 0.1%, while for cells in two adjacent trenches, the contribution is just under 0.3%. **c)** Cells situated directly beneath the mother cell also contribute intensity through bleedthrough effects. The top schematic illustrates the experiment with mixed labelled and unlabeled cells within a trench, and corresponding snapshots from the actual experiment are displayed below. The total contribution increases proportionally with the number of cells beneath the mother cell, but this effect plateaus rapidly (2.25%) as additional cells are introduced further away from the mother cell at the top.

### Supplementary Information 20: Multi-snapshot single molecule counting

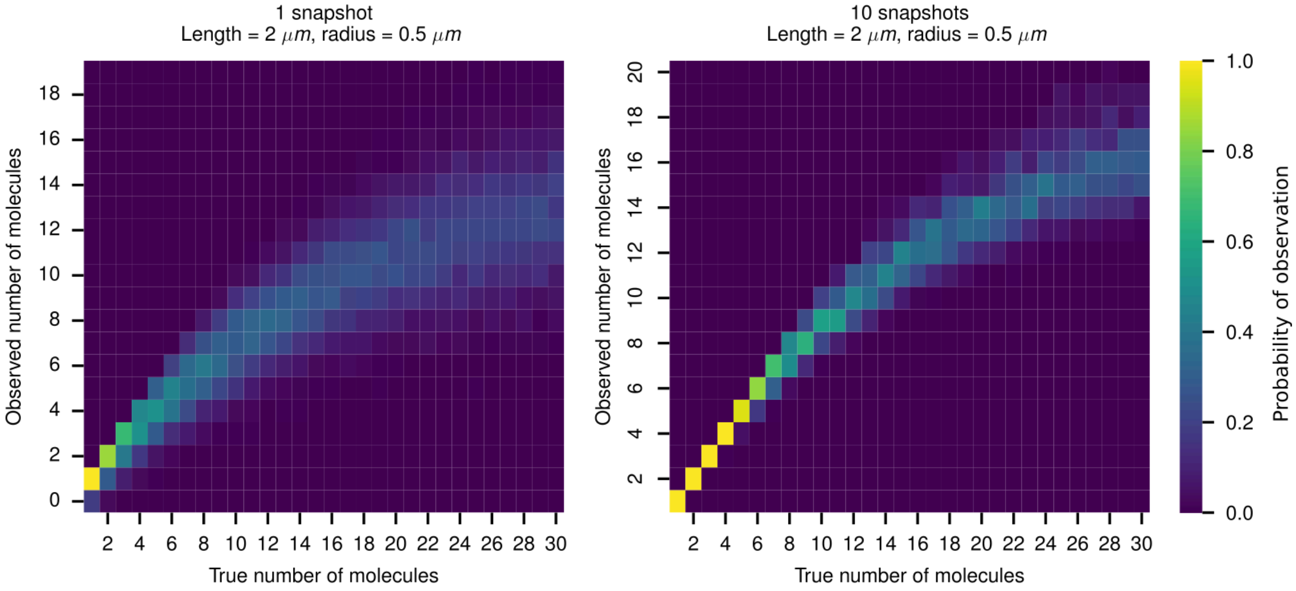


**Figure S20:** The heat maps show the probability of detecting a particular count as a function of input number of emitters (naive method). Picking the maximum count out of 10 snapshots leads to an improved performance compared to a single snapshot. Areas marked in green hatching indicate regions where probability of counting the number of molecules correctly is greater than 95%.

### Supplementary Information 21: Deriving a correction factor for depth dependent loss of molecules

Consider a cell containing a small number of single fluorescent molecules being imaged. We would like to model the probability of $N_{\mathrm{obs}}$ given that the cell has $N_{\mathrm{true}}$ molecules present. $N_{\mathrm{obs}}$ will often be less than $N_{\mathrm{true}}$ due to molecules either being too dim, out of focus, or in too close proximity to other molecules to be resolved. Let us consider the geometry of a single cell, which we model as a spherocylinder. The cross-sectional area of a cell in the $xy$ plane is given by

$$A(z) = 2L\sqrt{2rz - z^2} + \pi(2rz - z^2)$$

Where $L$ is the length of the mid-section of the cell (excluding the hemispherical caps), $r$ is the half-width of the cell and the radius of the hemispherical caps, and $z$ is the distance from the top of the cell. The first term is simply the area of the rectangular mid-section of the cell, and the second term is the area of the hemispherical end caps accounting for the depth of the cross-section.

A probability density function for finding molecules within a given depth of the cell can therefore be given by:

$$a(z) = \frac{A(z)}{\int_0^{2r} A(z)dz}$$

However, due to defocus dimming of molecules, some molecules have a chance of not being detected at various depths. We model the detection probability as a super-Gaussian:

$$D(z) = A \cdot e^{-\left(\frac{(z-z_0)^2}{2\sigma_z^2}\right)^P}$$

This super-Gaussian is a good fit to the empirically measured detection probability, which inherently takes into account the variability in the brightness of molecules. We modelled the empirical detection probability function by defocussing a single point source (modelled using the vectorial PSF model given by Aguet[2], fitted to our instrumental PSF). The point source was considered detectable if its SNR was greater than the 99th percentile of background PSNRs. The simulated molecule's photon counts were Poisson distributed such that at the focal plane the measured SNR was 8 on average.

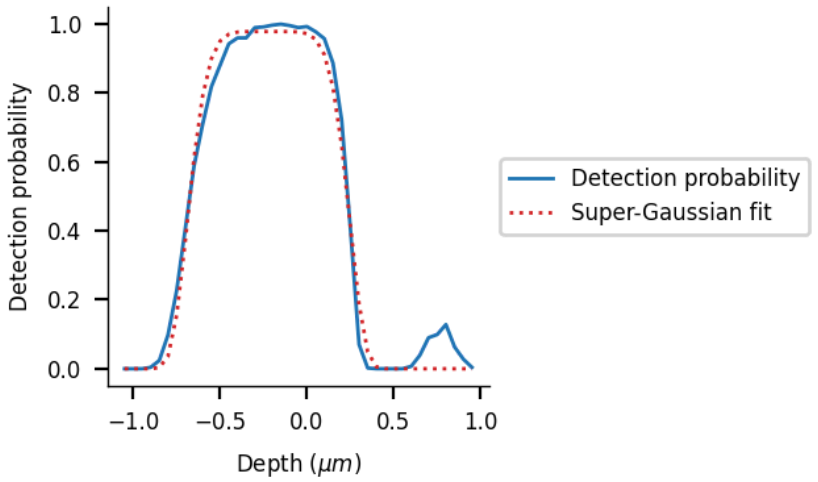


**Figure S21:** The detection probability estimated by the point spread function vs the super-Gaussian fit. The small hump between 0.5 and 1 micron is not modelled.

Therefore we can define the expected number of molecules observed as a function of the total number of molecules within the cell as a function of the density distribution of the molecules weighted by the detection probability:

$$\mathbb{E}[N_{\text{obs}}] = N_{\text{true}} \int_0^{2r} D(z)a(z)dz$$

(Noting that here, $D(z)$ is a shifted version for the purposes of integrating between $0$ and $2r$.

Therefore, using this scheme, $\frac{N_{\text{obs}}}{N_{\text{true}}}$ is a constant for a given cell geometry. This can be used to correct for the number of molecules lost to depth of field effects. This, however, will not correct for diffraction effects, and is therefore most useful at low molecular counts or when the depth of the sample is large. Additionally, we can use this information to adjust the experimental setup by defocussing in order to maximise the detection probability in a given geometry. Take for instance, a focal offset of $\delta z$, the maximum detection probability can be found by shifting the detection probability curve (assuming the objective moves, and not the microscope stage), and solving:

$$\frac{1}{N_{\text{true}}}\frac{d}{d\delta z}\mathbb{E}[N_{\text{obs}}](\delta z) = \int_0^{2r} \frac{\partial}{\partial \delta z} D(z - \delta z)a(z)dz = 0$$

Therefore given an optimal value of $\delta z$, and accurate modelling of the cell's geometry, one can correct for detection loss due to signal loss. It should be noted that correction factor applies to the average molecular count.

### Supplementary Information 22: Image simulation pipeline

Fluorescent image simulation consists of several steps. First, emitters are sampled within a 3D cell according to some distribution. In this paper we chose between uniformly distributed emitters within a spherocylindrical hull, or within the membrane/cell wall region of a cell. These 3D distributed emitters are then split into corresponding z-slices, and convolved with the corresponding z-slice of a point spread function. The convolved slices are then summed to create the final cell image, which is a projection of the 3D object, and is what is observed under the microscope. Image simulation is always performed initially at a very small pixel size (0.005 μm/pixel) in order to correctly sample all small features of a point spread function (when using synthetic PSFs), and also to accurately sample the roundness of a cell without pixelation. In the case where an iPSF is being used for convolution, it is upsampled using linear interpolation. For direct simulation to experimental comparison, images are then downsampled to the original experiment's pixel size.

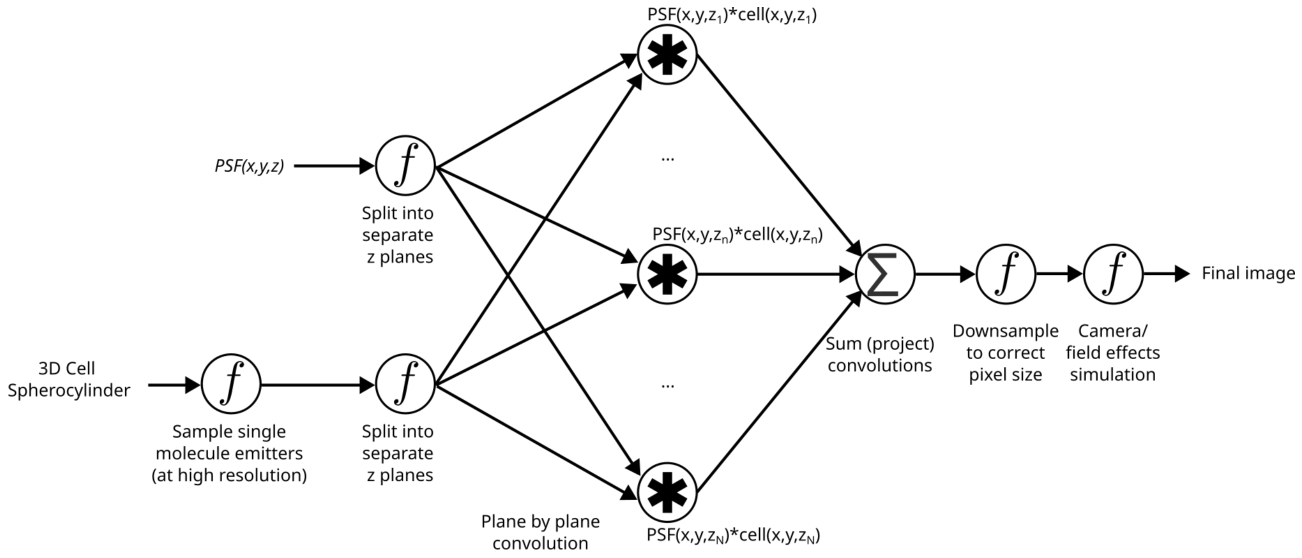


**Figure S22:** A block diagram representation of the basic image formation process, showing emitter distribution, plane by plane convolution, and projection.

#### Supplementary Information 23: Plane-by-plane convolution is necessary for accurate image simulation

Consider a 3D projected volume being imaged, made up on $N$ slices in the $z$ plane: $\{I_{z_1}, I_{z_2}, \ldots, I_{z_N}\}$. The resultant image ($C$) will be the sum of the convolution of each z plane of the PSF kernel with its corresponding image z plane:

$$C = \sum_{i=1}^{N} (I_{z_i} * K_{z_i})$$

This result is different from the naive method of convolution (which would result in a false image, $C_f$ ), which would involve convolving the sum the kernel planes with the sum of the volume planes

$$C_f = \left( \sum_{i=1}^{N} I_{z_i} \right) * \left( \sum_{i=1}^{N} K_{z_i} \right)$$

From the additivity of convolution we can show that this is equivalent to:

$$\left( \sum_{i=1}^{N} I_{z_i} \right) * \left( \sum_{i=1}^{N} K_{z_i} \right) \\ = \sum_{i=1}^{N} \sum_{i=1}^{N} (I_{z_i} * K_{z_i})$$

This is a double sum that convolves each image with every kernel and sums all these convolutions. This is fundamentally different from the first expression, where each image is convolved with its respective unique kernel and only these convolutions are summed.

A similar principle holds for deconvolution. Since we only have access to the final projected image $C$, information about the convolution process of each layer is essentially lost, making correct deconvolution impossible.

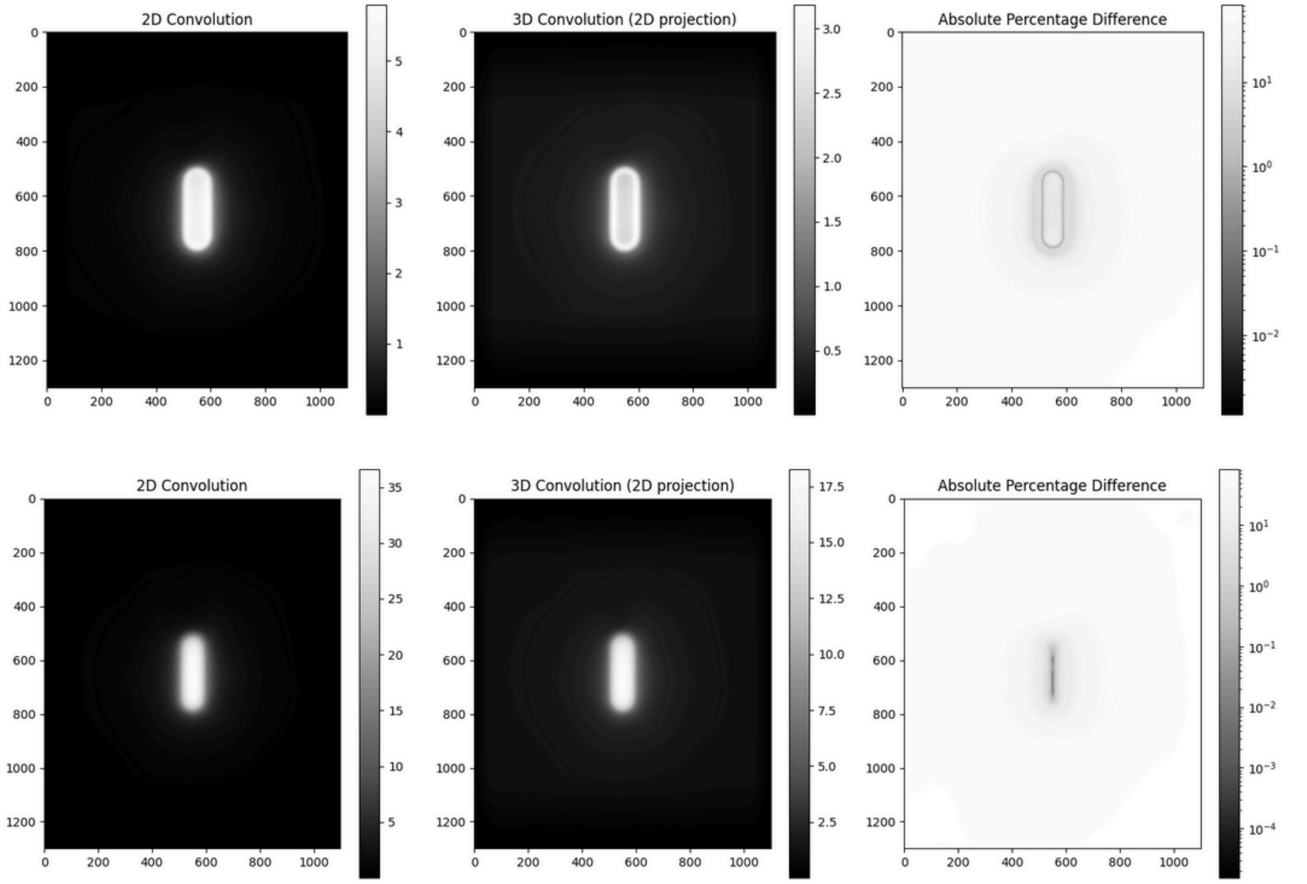


**Figure S23**: Showing the difference between naive 2D convolution vs 3D convolution + projection in both membrane and cytoplasmically fluorescent cells. The resultant images can be markedly different. Only correct deconvolution of the first image would be possible, as it is the direct inverse of the operation which created it. Deconvolution of the projected image is impossible since information is lost in the layer summation process. Thus single cell size estimation cannot be corrected for by deconvolution.

#### Supplementary Information 24: Segmentation of microcolonies for extracting single-cell fluorescence data

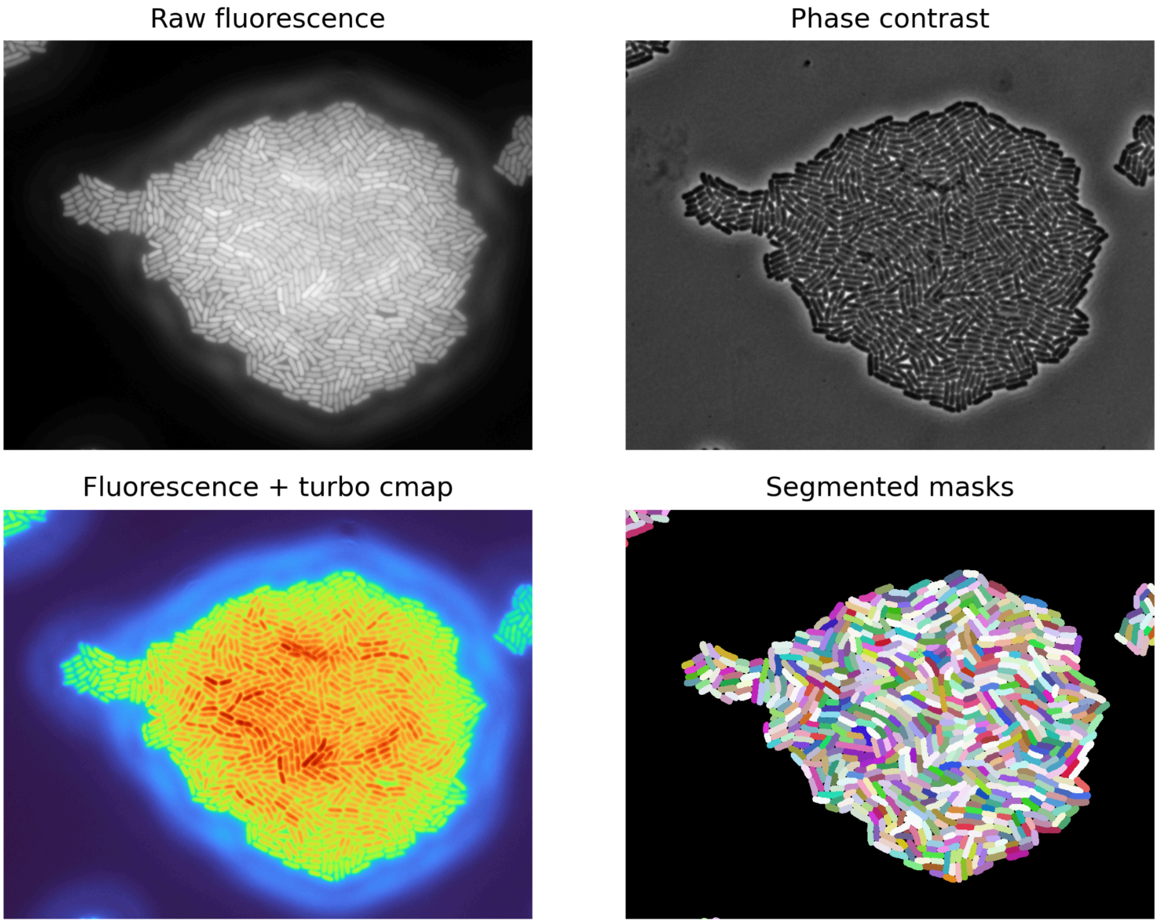


**Figure S24:** Example fluorescence and phase-contrast images of a microcolony (top panels) and the corresponding colour map of fluorescence intensities and single-cell masks from phase-contrast image segmentation are shown (bottom panels).


1. Hanser, B. M., Gustafsson, M. G. L., Agard, D. A. & Sedat, J. W. Phase-retrieved pupil functions in wide-field fluorescence microscopy. *J. Microsc.* **216**, 32–48 (2004).
2. Aguet, F., Geissbühler, S., Märki, I., Lasser, T. & Unser, M. Super-resolution orientation estimation and localization of fluorescent dipoles using 3-D steerable filters. *Opt. Express* **17**, 6829–6848 (2009).
